# Single-cell RNA sequencing unveils intestinal eosinophil development and specialization

**DOI:** 10.1101/2021.10.27.466053

**Authors:** Alessandra Gurtner, Costanza Borrelli, Nicolás G. Núñez, Ignacio Gonzalez-Perez, Daniel Crepaz, Kristina Handler, Tomas Valenta, Konrad Basler, Burkhard Becher, Andreas E. Moor, Isabelle C. Arnold

## Abstract

Eosinophils are an integral part of the gastrointestinal (GI) immune system that contribute to homeostatic and inflammatory processes. Here, we investigated the existence of functional subsets that carry out specialized tasks in health and disease. We used single-cell transcriptomics and high-dimensional flow cytometry to delineate murine eosinophil subpopulations and their ontogenetic relationship in the steady state and during infection and inflammation. Profiling of eosinophils from bone marrow, blood, spleen and several GI tissues revealed five distinct subsets representing consecutive developmental and maturation stages across organs, each controlled by a specific set of transcription factors. Furthermore, we discovered a highly adapted PD-L1^+^CD80^+^ eosinophil subset in the GI tract, characterized by its immune regulatory properties, bactericidal activity upon challenge infection and tissue-protective function during inflammation. Our data provide a framework for the characterization of eosinophil subsets in GI diseases and highlight their crucial contribution to homeostasis, immune regulation and host defense.

## Introduction

Eosinophils arise in the bone marrow (BM) from lineage-committed progenitors (Iwasaki et al., 2005) and are released into the blood stream as mature cells. Eosinophils then rapidly migrate to their target tissues, where their lifespan is extended by endogenous cytokines. Under homeostatic conditions, small numbers of eosinophils reside in several tissues, including the thymus, uterus, lung and gastrointestinal (GI) tract (Marichal et al., 2017). Eosinophil accumulation in peripheral organs is typical of disease states such as allergic airway inflammation, atopic dermatitis and eosinophilic esophagitis (Blanchard et al., 2006; Humbles et al., 2004; Jenerowicz et al., 2007) and is also a common feature of inflammatory bowel diseases (IBD) (Raab et al., 1998).

The largest population of tissue-resident eosinophils is found in the healthy GI lamina propria (LP) of mice and humans, where they contribute to various intestinal homeostatic processes, including epithelial barrier preservation, tissue architecture support, immune cell population maintenance as well as regulation of local immune responses (Chu et al., 2014; Jung et al., 2015; K. Shah, 2021; Sugawara et al., 2016). During GI inflammation, eosinophil numbers dramatically increase, with evidence of local degranulation and DNA extracellular trap (EETs) release (Yousefi et al., 2008). However, eosinophils are not systematic drivers of pathology and their function during intestinal inflammation is unclear (Alhmoud et al., 2020).

Eosinophils exhibit considerable phenotypical, morphological and functional diversity within tissues (Mesnil et al., 2016; Xenakis et al., 2018), indicating the existence of specialized subset with tissue-protective or pro-inflammatory functions (Abdala-Valencia et al., 2018; Masterson et al., 2021). Thus, delineation of eosinophil subpopulations is required to develop refined targeting strategies harnessing eosinophil pathological activities. However, the presence of transcriptionally defined eosinophil subsets and their ontogenetic relationship has remained unexplored to date due to technical challenges preventing their transcriptomic interrogation.

Here, we developed a method that minimizes shear stress, cell degranulation and transcript degradation to resolve eosinophil transcriptional heterogeneity at the singlecell level. We aimed to comprehensively analyze eosinophil profiles from their differentiation in the BM to their tissue of residence – at steady state and during inflammation – in order to examine the existence of eosinophil subsets, understand their developmental trajectory and define their functional contribution to intestinal homeostatic and inflammatory processes.

## Results

### Single-cell transcriptomic profiling reveals five distinct eosinophil subpopulations across tissues

To investigate eosinophil transcriptional profiles throughout development, we analyzed Siglec F^+^ cells magnetically isolated from the blood, spleen, stomach, small intestine (SI) and colon of *Il5*-tg mice (Figure S1A). To sample the entire eosinophil lineage, whole BM extracts were included. Cells from different organs were barcoded and multiplexed to minimize batch effects, and were sequenced to saturation on the BD Rhapsody platform (Figures 1A and S1B). Following removal of contaminant cell clusters not belonging to the eosinophilic lineage (Figure S1C,D), we identified 7,796 single cells expressing the key eosinophils markers *Siglecf*, *Il5ra*, *Ccr3* and *Epx* (Figure S1E). Louvain clustering revealed five main eosinophil subpopulations (Figures 1B and S1F), two of which were present primarily in the BM (termed progenitors and immature eosinophils) and one was enriched in the blood (circulating eosinophils). The remaining two subpopulations (basal and active eosinophils) predominantly populated peripheral tissues, but in varying proportions across organs (Figure 1C,D).

**Figure 1:**
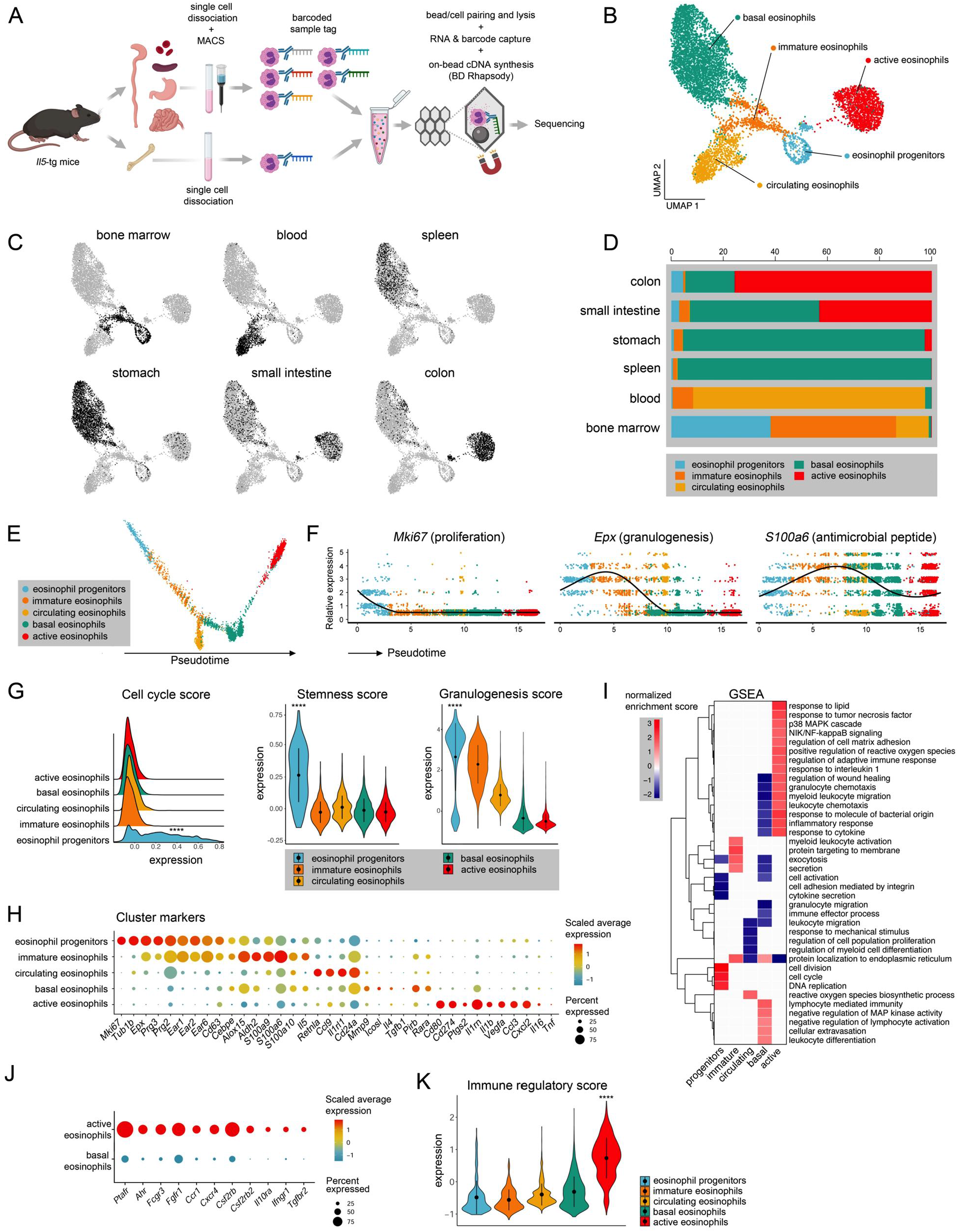
scRNA-seq reveals five distinct eosinophil subpopulations. **A)** Experimental workflow. **B)** UMAP of eosinophil transcriptomes (n = 3, *Il5*-tg mice). **C)** Organ-derived eosinophil distribution. **D)** Eosinophil subset distribution across organs (% of total eosinophils). **E)** Eosinophil development trajectory. **F)** Expression of *mKi67*, ***E**px* and ***S100a6*** over pseudotime. **G)** Left: cell cycle score. Middle: stemness score. Right: granulogenesis score. **H)** Selected marker gene expression. **I)** Significantly enriched (adjusted P < 0.05) GSEA terms across clusters. **J)** Receptor gene expression. **K)** Immune-regulatory score. B,E,F) Dots represent single cells colored by cluster identity. G, K) Genes used for scores and signatures are listed in Table S2. Data represent the mean ± SD. ****P < 2.2×10^−16^ (two-sided Wilcoxon test). See also Figure S1.

Ordering of eosinophil subsets along pseudotime placed eosinophil progenitors at the origin of the differentiation trajectory, with immature, circulating, basal and active eosinophils representing consecutive post-mitotic subpopulations ordered along a developmental continuum (Figure 1E). The identity of eosinophil progenitors was evidenced by strong expression of proliferation, cell cycle control and core stemness genes (Koeva et al., 2011) (Figures 1F,G and S1G). As previously reported (Bouffi et al., 2015), eosinophil progenitors were found to undergo active granulogenesis (Figure 1G), with robust expression of granule proteins genes such as *Epx*, *Prg2* and *Ear1* (Figure 1H). As eosinophils progressed to the immature stage, stemness and proliferation genes were downregulated, while genes encoding antimicrobial peptides (*S100a6,9,10*) were upregulated (Figures 1F–H and S1G). The significant enrichment of gene sets related to protein targeting to the membrane, secretion and exocytosis further indicated sustained granule biogenesis in this subset (Figure 1I) and were downregulated as eosinophils matured and entered the circulation. Rather surprisingly, circulating eosinophils consisted of a transcriptionally separate cluster, characterized by high expression of the cell adhesion molecule *Cd24a* and the cysteine-rich secreted protein *Retnla*, as well as upregulation of genes involved in reactive oxygen species biosynthesis (Figure 1H, I).

Following migration to the GI tract, eosinophils bifurcated into two transcriptionally distinct subpopulations, termed hereafter “basal” and “active” eosinophils (Figure 1B). Basal eosinophils were found in all organs of the GI tract, but also constituted the predominant subset in the spleen (Figure 1C,D). They expressed cytokines such as *Il4*, *Il13* and *Tgfb1,* as well as the co-stimulatory molecule *Icosl* and the extracellular matrix (ECM)-degrading enzyme *Mmp9* (Figure 1H). Basal eosinophils further upregulated the inhibitory receptor *Pirb* and the alpha subunit of the retinoic acid receptor (*Rara*; Figure 1H), signaling from which has been linked to enhanced eosinophil survival (Ben Baruch-Morgenstern et al., 2014; Ueki et al., 2008).

In contrast, active eosinophils were predominantly enriched in the colon and SI (Figures 1C,D and S1H). Their emergence at the end of the eosinophil differentiation trajectory indicated their terminal maturation state (Figure 1E). This subset was characterized by robust synthesis of cytokines (*Il16*, *Tnf, Il1b* and its receptor antagonist *Il1rn*), chemokines (*Ccl3*, *Cxcl2*), growth factors (*Vegfa*) and the prostaglandin synthetase Cox-2 (*Ptgs2;* Figure 1H). In addition, active eosinophils upregulated multiple receptor genes, including cytokine and chemokine receptors (*Il10ra*, *Csf2rb*, *Tgfbr2*, *Ccr1*, *Cxcr4*), the platelet-activating factor receptor (*Ptafr*) and the aryl hydrocarbon receptor (*Ahr*; Figure 1J). Active eosinophils further upregulated gene sets involved in immune modulation (Figure 1I). Indeed, expression of the co-stimulatory molecules *Cd80*, *Cd274* and *Cd9* was significantly higher than in basal eosinophils (Figure 1K) and peaked at the end of the differentiation trajectory (Figure 2A). The expression of genes encoding multiple bioactive factors, surface receptors and regulatory molecules characteristic of active eosinophils suggests this subpopulation exerts critical tissue-specific functions in the GI tract. We therefore focused our attention on this subset and examined its phenotypic and functional characteristics in relation to its basal counterpart.

**Figure 2:**
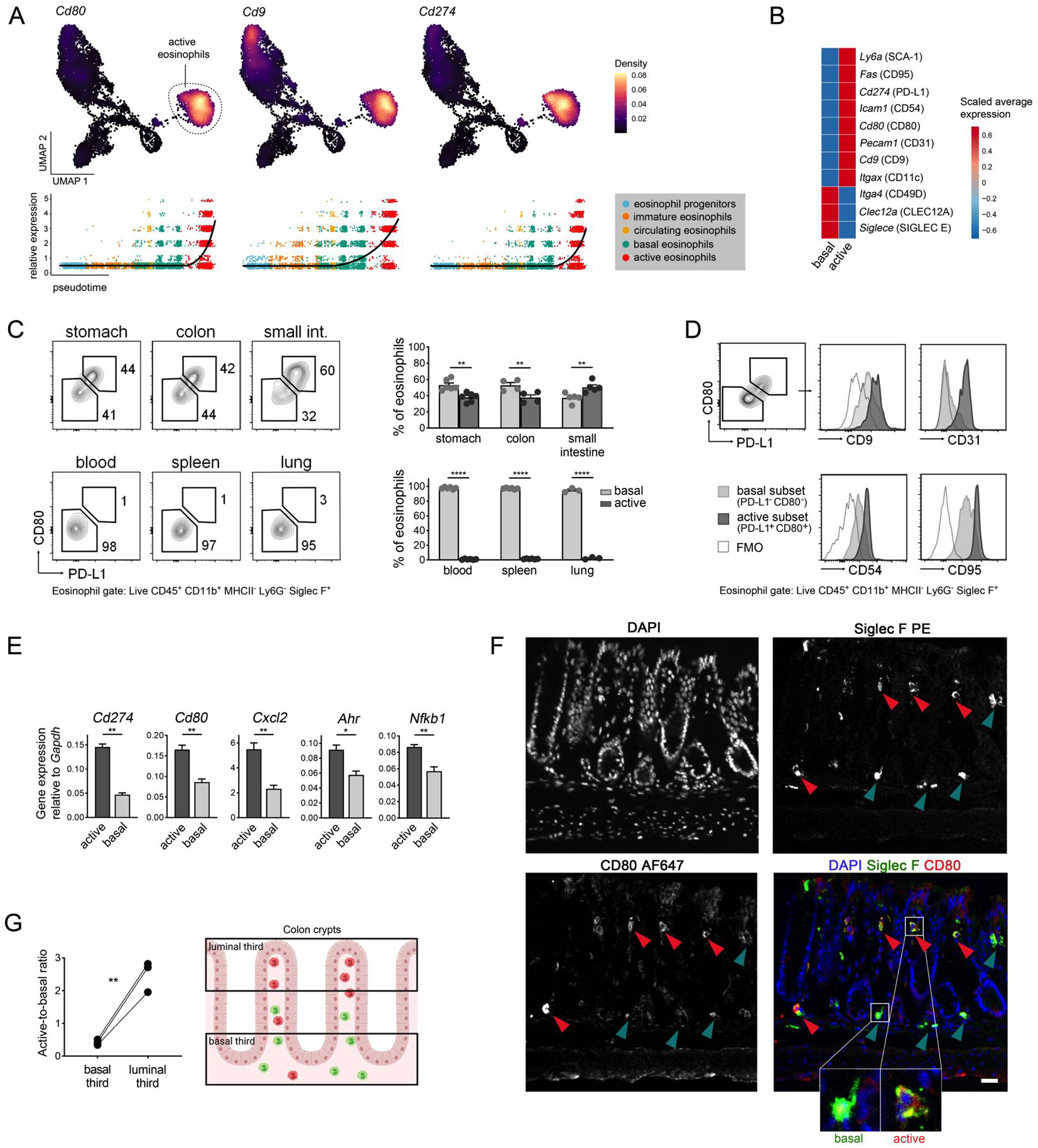
PD-L1 and CD80 expression define active eosinophils in the GI tract. **A)** Top: UMAP of *Cd80*, *Cd9* and *Cd274* expression. Bottom: expression over pseudotime. Dots represent single cells. **B)** GI surface marker expression. **C)** Left: representative flow cytometry plots of active (CD80^+^ PD-L1^+^) and basal (CD80^−^ PD-L1^−^) eosinophils. Right: frequencies of active and basal eosinophils (n = 4-6, B6J mice). Data represent the mean ± SEM. **D)** Mean fluorescence intensity (MFI) of CD9, CD31, CD54 and CD95 across colonic eosinophil subsets. FMO: fluorescence minus one. **E)** Gene expression normalized to *Gapdh* measured by qRT-PCR of active vs. basal eosinophils sorted from the SI of *Il5*-tg mice (n = 4). Data represent the mean ± SEM. **F)** Representative images of eosinophil immunofluorescent staining for Siglec F and CD80 in the colon of B6J mice. Arrows mark Siglec F^+^ CD80^+^ active eosinophils (red) and Siglec F^+^ CD80^−^ basal eosinophils (green). Dotted line delimits the epithelial layer. Nuclei are stained with DAPI. Scale bar, 20 *μ*m. **G)** Active-to-basal eosinophil ratio in luminal (upper) third vs. basal (lower) third colonic crypts of B6J mice (n = 3). **P < 0.01 (unpaired t-test). C,E) *P < 0.05; **P < 0.01; ****P < 0.0001 (Mann-Whitney test). See also Figure S2.

### PD-L1 and CD80 expression define active eosinophils in the GI tract

To characterize active eosinophils within the GI tract, we first defined a set of surface markers suitable for their discrimination by flow cytometry. To this end, we extracted surface protein genes from each cluster (Figure S2A) and selected those upregulated in the GI tract (Figure 2B). The active subset was best defined by co-expression of *Cd274* (PD-L1) and *Cd80,* at both the transcript (Figures 2A and S2B) and protein levels (Figure 2C). As predicted by our transcriptomic data, PD-L1^+^ CD80^+^ eosinophils were restricted to the GI tract and expressed higher levels of CD9, CD31, CD54 and CD95 than their PD-L1^−^ CD80^−^ basal counterparts (Figure 2D), as well as higher intensity of the activation marker Siglec F (Figure S2C). Analysis of colonic eosinophils sorted by flow cytometry further confirmed the expression of subset-specific transcripts (Figure 2E). Of note, protein products of genes expressed exclusively in the basal subset (*Siglece*, *Itga4*, *Clec12a*) were also expressed in active eosinophils (Figure S2D), possibly reflecting their low turnover rate. Interestingly, both Siglec F^+^ CD80^−^ basal and Siglec F^+^ CD80^+^ active subsets were readily found in the colonic LP of mice by immunofluorescent staining (Figure 2F) and differed in their mucosal localization (Figure 2G). Active eosinophils were found significantly closer to the luminal extremity of the mucosa (luminal third), while basal eosinophils were retained near the basement membrane (basal third). Together, we conclude that PD-L1 and CD80 represent *bona fide* RNA and protein markers of active eosinophils that can be used for their identification *in vivo*.

### Active eosinophils undergo regional tissue adaptation and can be induced experimentally *in vitro* and *in vivo*

Active eosinophils express multiple receptors (Figure 1J), suggesting responsiveness to tissue-specific cues. To assess the adaptability of active and basal eosinophils to distinct GI niches, we compared their profiles across the diverse organs of the GI tract. Interestingly, basal eosinophils displayed a homogeneous transcriptome irrespective of their GI compartment (Figures 3A and S3A) and did not significantly differ from those residing in the spleen (Figure S3B). In contrast, active eosinophils exhibited strong organ-specific gene signatures, with the greatest diversity found among colonic eosinophils (Figures 3A and S3A). In the colon, active eosinophils exhibited enhanced expression of genes such as *Itgal, Serpinb2*, *Tgm2* and *Thbs1*, which are involved in interactions with, and remodeling of ECM components. In the SI, they upregulated genes such as *Cd300lf* and the short-chain fatty acids receptor *Ffar2*, signaling from which was recently associated with reduced eosinophil viability (Theiler et al., 2019).

**Figure 3:**
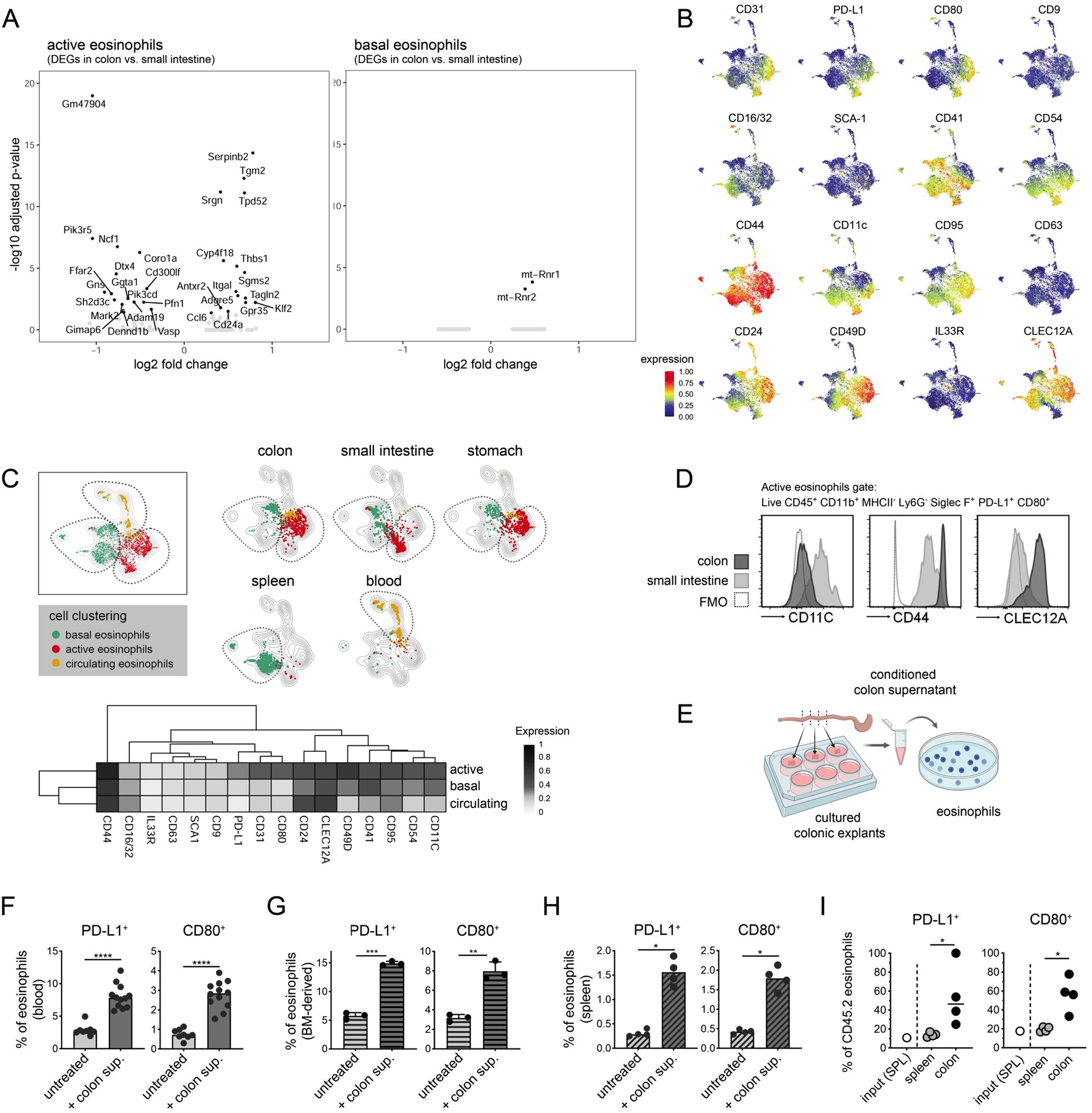
Active eosinophils undergo regional tissue adaptation and can be induced experimentally *in vitro* and *in vivo*. **A)** Significant DEGs (black dots, average log fold-change > 0.5, adjusted P < 0.05) between colon and SI within the active (left) and basal (right) clusters. **B)** UMAP showing the normalized expression intensity of the 16 surface markers on eosinophils (n = 3-4, B6J mice). **C)** Top: UMAP showing the manual annotation of FlowSOM clusters. Bottom: Heatmap of median surface marker expression across subsets. **D)** MFI of CD11c, CD44 and CLEC12A in active eosinophils shown in C. **E)** Workflow of *in vitro* conditioning. **F-H)** Frequencies of PD-L1^+^ and CD80^+^ blood eosinophils (F), BM-derived eosinophils (G) and splenic eosinophils (H) after conditioning with colon supernatant (n = 3-4, isolated from *Il5*-tg (F, H) or B6J (G) mice). In F), data are pooled from three independent experiments. Data represent the mean ± SEM. **I)** Frequencies of PD-L1^+^ and CD80^+^ eosinophils among adoptively transferred CD45.2^+^ eosinophils isolated from the spleen and colon after 42 h. Input cell (splenic eosinophils) frequency is shown as a reference (n = 4, CD45.1 mice). Medians are shown. *P < 0.05; **P < 0.01; ***P < 0.001; ****P < 0.0001 (Mann-Whitney test). See also Figure S3.

To further clarify GI eosinophil phenotypes, we characterized blood, spleen and GI eosinophils of wild-type (WT) mice by high-parametric spectral flow cytometry using 16 independent markers. As predicted from our RNA dataset, the expression patterns of PD-L1, CD80 and CD31 predominantly overlapped and were exclusively expressed by eosinophils of GI tissues (Figure 3B,C). FlowSOM analysis confirmed the presence of three major eosinophil subpopulations (basal, active and circulating) based on their surface marker profiles (Figure 3C). Within the active cluster, SI eosinophils adopted a phenotype that is distinct from their colonic counterparts and were characterized as CD11c^hi^ CD44^lo^ CLEC12A^lo^ (Figures 3D and S3D,E).

We next assessed eosinophil maturation into the active subset experimentally. Purified blood or BM-derived (BM-eos) eosinophils were conditioned with the supernatant of cultured colonic explants (colon supernatant, Figure 3E), resulting in enhanced PD-L1^+^ and CD80^+^ surface expression (Figure 3F,G). Similarly, the conditioning of splenic (basal) eosinophils induced their differentiation into the active phenotype (Figure 3H). The *in vivo* adoptive transfer of CD45.2 splenic (basal) eosinophils into CD45.1 murine hosts further resulted in their migration to the colon, where they showed evidence of *in situ* maturation (Figure 3I). In contrast, CD45.2 eosinophils recovered from the spleen retained low expression of PD-L1^+^ and CD80^+^ (Figure S3F). These observations suggest that basal-to-active eosinophil conversion is possible upon appropriate stimulation and is driven locally by the intestinal milieu.

### Active eosinophil differentiation is driven by NF-κB signaling and relies on bacterial cues

We next examined the key factors and pathways underlying active eosinophil differentiation. Single-cell regulatory network inference and clustering (SCENIC) analysis revealed highly diverse regulon activities across clusters and a non-overlapping transcription factor (TF) profile specific to each subset (Figures 4A and S4A). In basal eosinophils, increased activity of the TF Meis1 was related to the expression of signatures genes such as *Mmp9*, *Tgfb1*, *Ccr3* and *Itga4* (CD49d). Stat6 regulon activity was elevated in this subset (Figure S4A), while Stat3 regulon activity prevailed in active eosinophils (Figure 4A,B). Additional TFs selectively predicted to govern the transcriptional landscape of active eosinophils included Ahr, Irf5 and HiF1-α (Figure 4A,B). Active eosinophils further showed striking upregulation of several NF-κB-related regulons, including Rela, Relb, Nfkb1 and Nfkb2 (Figure 4A,B), indicating robust activation of this pathway. Accordingly, NF-κB target genes and pathway components were selectively upregulated in active eosinophils (Figure 4C) and were expressed at significantly higher levels in colonic eosinophils compared with their blood and splenic counterparts (Figure 4D). Notably, NF-κB TFs were predicted to regulate expression of *Cd274*, *Cd80* and *Cd9*. Indeed, the co-localization of phosphorylated NF-κB p65 (p-NF-κB p65) with CD80^+^, but not CD80^−^ eosinophils in the colonic LP indicates that canonical NF-κB signaling is selectively induced in active eosinophils (Figure 4E,F). PROGENy analysis of pathway activity confirmed an NF-κB transcriptional signature in active eosinophils and further indicated high TNF-α signaling, an upstream mediator of the NF-κB pathway (Figure S4B). However, PD-L1^+^ CD80^+^ active eosinophil frequencies were unaltered in TNF receptor 1-deficient mice (TNFR1^-/-^) (Figure S4C). Similarly, *in vivo* neutralization of IL-5 and GM-CSF, known to signal partly via the NF-κB pathway, failed to abrogate active eosinophil differentiation (Figure S4D), indicating that these cytokines do not contribute to active eosinophil maturation.

**Figure 4:**
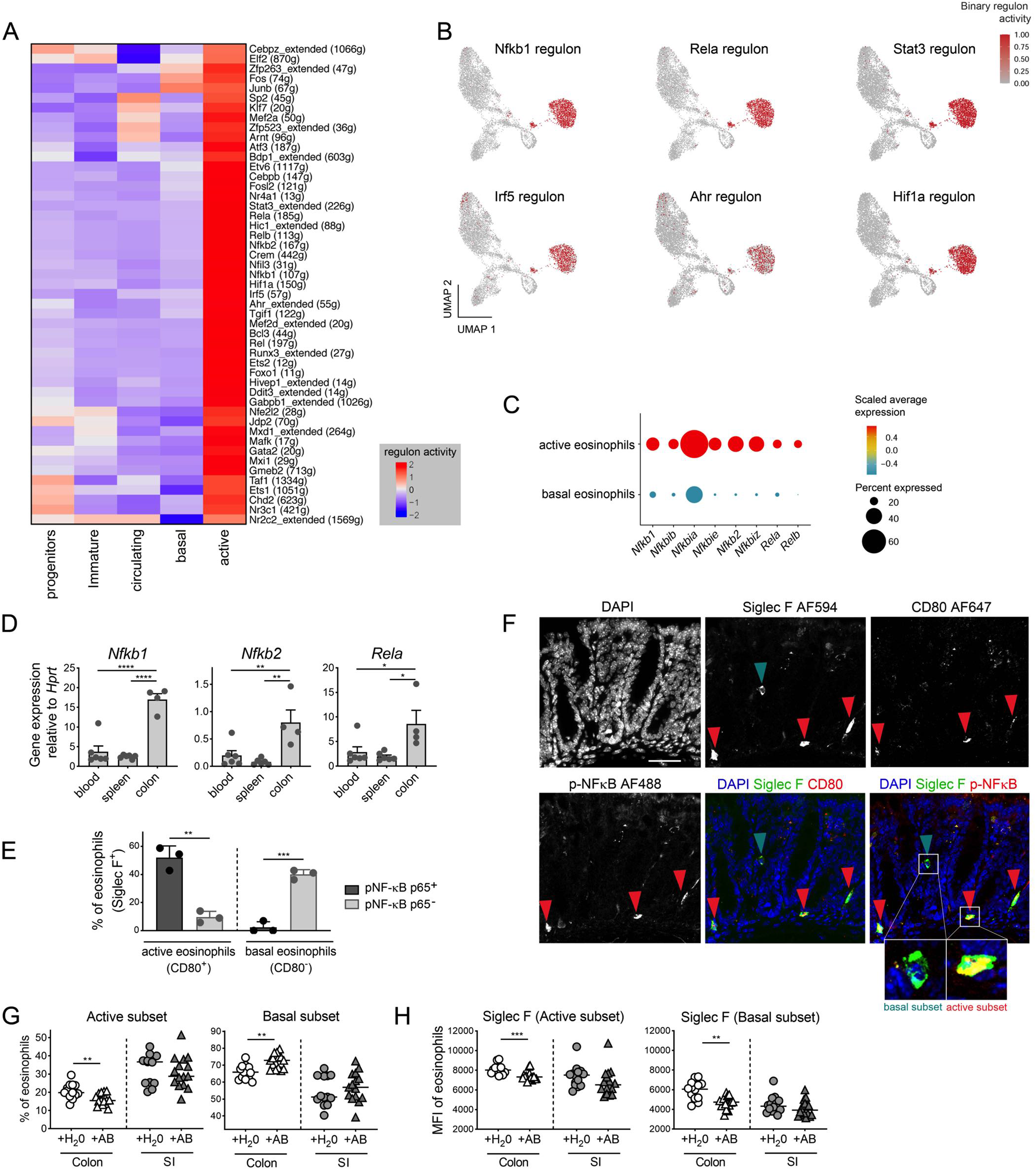
Active eosinophil differentiation is driven by NF-κB signaling and relies on bacterial cues. **A)** Regulon activity across clusters. Active subset-specific regulons are shown. **B)** UMAP of Nfkb1, Rela, Stat3, Irf5, Ahr and Hif1a regulons. Cells are colored by binary regulon activity. **C)** Expression of NF-κB signaling components. **D)** Gene expression relative to Hprt measured by qRT-PCR of sorted eosinophils (n = 4-6, *Il5*-tg mice). Data represent the mean ± SEM. *P < 0.05; **P < 0.01; ****P < 0.0001 (One-way ANOVA). **E-F)** Quantification (E) and representative image (F) of pNF-κB p65 immunofluorescent staining in colonic eosinophils. Arrows mark active (CD80^+^, red) and basal (CD80^−^, green) Siglec F^+^ eosinophils. Three fields of view are quantified from three biological replicates (B6J mice). Nuclei are stained with DAPI. Scale bar, 20 *μ*m. **P < 0.01; ***P < 0.001 (unpaired t-test). **G)** Frequencies of active and basal eosinophils treated with antibiotics relative to controls (n = 14-15, B6J mice). Data are pooled from two independent experiments. **H)** Activation status of intestinal eosinophils shown in G assessed by MFI of Siglec F. G,H) Medians are shown. **P < 0.01; ***P < 0.001 (Mann-Whitney test). See also Figure S4.

The selective presence of active eosinophils within the GI tract and the upregulation of gene modules involved in response to bacterial stimulation (Figure 1I) suggests their differentiation is driven by microbiota-derived cues. We therefore evaluated the frequency of active eosinophils in mice treated with broad-spectrum antibiotics to deplete commensal bacteria. Notably, the proportion of PD-L1^+^ CD80^+^ active eosinophils in the colon was significantly reduced after 10 days of treatment (Figure 4G) and eosinophil activation status decreased (as defined by lower Siglec F intensity, Figure 4H). In contrast, the basal-to-active eosinophil ratio within the SI was only minimally affected (Figure 4G). Eosinophil maturation into the active subset *in vivo* and after *in vitro* conditioning was further not affected by TLR-2, TLR-4 or Myd88 deficiency (Figure S4E, F), suggesting independence of major bacterial recognition pathways.

### Challenge infection induces a compositional shift toward the active eosinophil cluster

Eosinophils populate the healthy GI tract as either basal or active population. However, it is unclear how local inflammation would affect eosinophil subset dynamics. We therefore investigated the subset-specific transcriptional responses to infection with two different GI pathogens – *Helicobacter pylori* (*H. pylori*) in the stomach or *Citrobacter rodentium (C. rod)* in the colon. We profiled eosinophils from the BM, blood and colon of *C. rod*-infected *Il5*-tg mice, and from the stomach of *H. pylori-*infected mice (Figure 5A). The single-cell profiles were integrated in the steady state transcriptional embedding and mapped with high confidence to the existing clusters (Figure S5A). Of note, merging the steady state and challenge datasets did not reveal novel infection-specific clusters (Figure S5B). Instead, both pathogens induced a strong compositional shift among eosinophil subsets at the sites of infection. Indeed, the proportion of active eosinophils strongly increased at the expense of their basal counterparts in infected organs (Figure 5B,C). At the protein level, eosinophils expressed higher intensities of PD-L1 and CD80 after bacterial challenge (Figure 5D) and PD-L1^+^ CD80^+^ active eosinophil frequencies increased markedly in the colonic and gastric LP of B6J mice relative to their basal counterparts (Figure 5E). Eosinophil quantification revealed a substantial accumulation of active eosinophils at infection sites, while basal eosinophil numbers remained unchanged (Figure S5C). High-parametric flow cytometry further confirmed a phenotypic change among eosinophil populations during bacterial challenge, characterized by higher expression of active eosinophil markers (Figures 5F,G and S5D,E).

**Figure 5:**
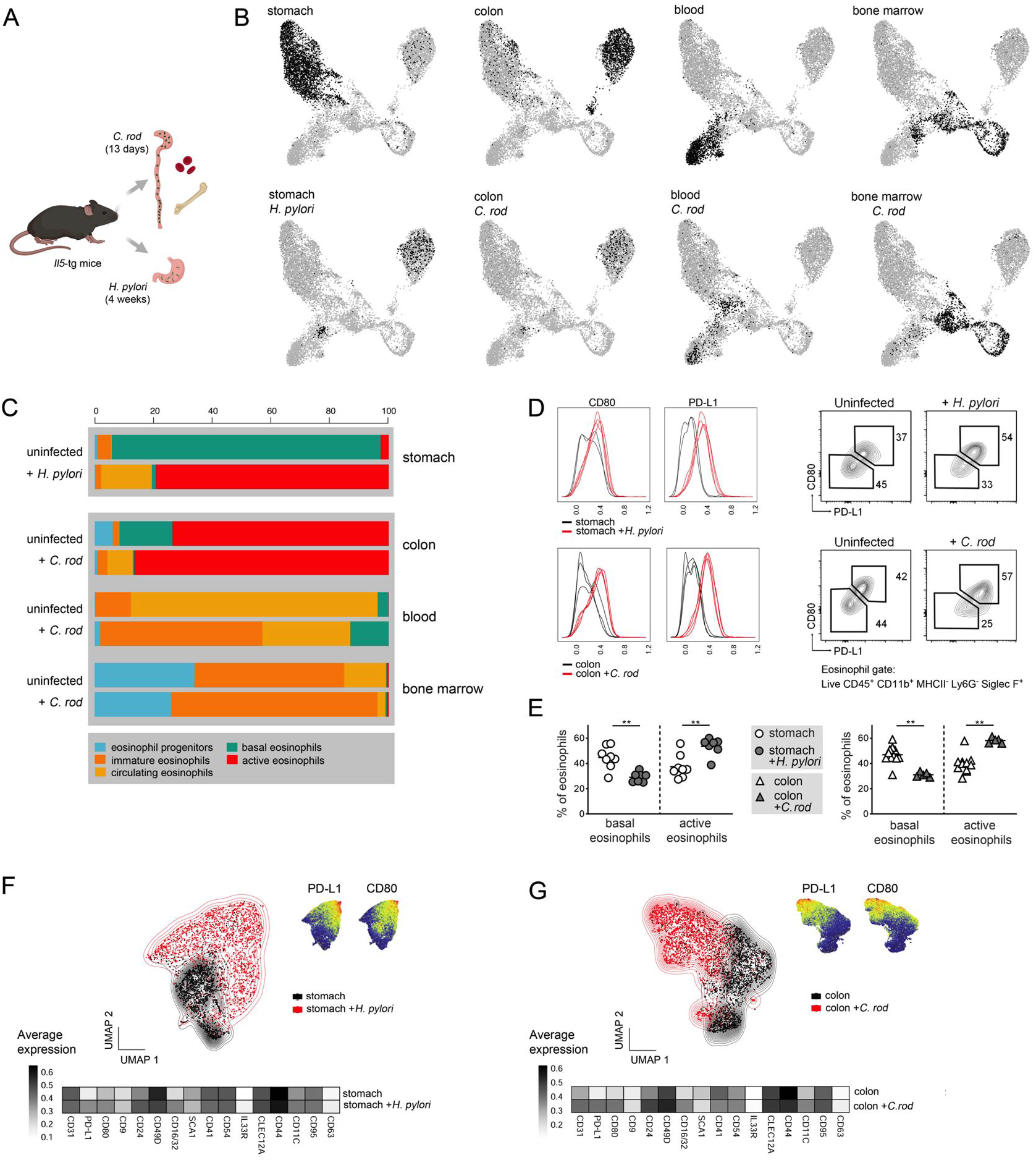
Challenge infection drives a compositional shift toward the active eosinophil cluster. **A)** Schematic diagram of bacterial challenge conditions. **B)** Organ-specific distribution of eosinophils (black dots) at steady state (top) and during bacterial infection (bottom; n = 3-4, *Il5*-tg mice). **C)** Percentage of eosinophil subsets across organs at steady state and after infection. **D)** Left: expression of CD80 and PD-L1 in eosinophils in uninfected (black) and infected (red) mice. Right: representative flow cytometry plots of the active (CD80^+^ PD-L1^+^) and basal (CD80^−^ PD-L1^−^) subsets. Numbers indicate the percentage of total eosinophils. Top: *H. pylori* in stomach, bottom: ***C.** rod* in colon. **E)** Percentage of active and basal eosinophils in *H. pylori*-infected stomach (left) and ***C.** rod*-infected colon (right) relative to uninfected controls (n = 7-10, B6J mice). Data are pooled from two independent experiments. Medians are shown. **P < 0.01 (Mann-Whitney test). **F-G)** UMAP of high-dimensional flow cytometric analysis of eosinophils in *H. pylori*-infected stomach (F) or ***C.** rod*-infected colon (G) relative to uninfected organs. Top right: UMAP showing PD-L1 and CD80 normalized expression intensity. Bottom: Heatmap of median surface marker expression across conditions (n = 4, B6J mice). See also Figure S5.

Infection also induced the accumulation of circulating eosinophils into the colon and stomach (Figure 5C). This population upregulated signature genes of active eosinophils such as *Tgm2*, *Vegfa*, *Nfkb2*, *Cd274*, *Csf2rb* and *Ahr* (Figure S5F), suggesting that it may convert directly into the active state by bypassing the basal maturation stage. Indeed, trajectory inference placed active eosinophils as originating directly from immature eosinophils during infection (Figure S5G). Eosinophil populations in the blood and BM of *C. rod-*infected animals also displayed a striking compositional shift compared to the steady state, with a relative expansion of immature eosinophils (Figure 5C). Hence, during infection, core eosinophil populations are maintained, but their proportions across organs vary to maximize active eosinophil production at sites of infection.

### Challenge infection promotes antimicrobial gene programs in active eosinophils

Expansion of active eosinophils during *H. pylori* and *C. rod* infection suggests a corresponding increase in their previously described regulatory and bactericidal activity (Arnold et al., 2018). We therefore examined the transcriptional changes occurring in this subset upon bacterial challenge. Both pathogens induced deregulation of multiple genes that were either pathogen-specific or shared among infection settings (Figures 6A and S6A). Active eosinophils apoptosis score decreased (Figure 6B) and biological processes related to cell activation, secretion/exocytosis and antimicrobial responses were enriched (Figure S6B). Indeed, both pathogens elicited a robust antimicrobial gene program, characterized by the expression of cathelicidin (*Camp*), lipocalin 2 (*Lcn2*), lactotransferrin (*Ltf*), calprotectin (a heterodimer of the *S100a8* and *S100a9* gene products) and lysozyme (*Lyz2*) (Figure 6C,D). The increased synthesis of bactericidal molecules was paralleled by upregulation of genes encoding eosinophil cytotoxic granule proteins (*Prg2*, *Epx*, *Ear1*, *Ear2*, *Ear6*; Figure 6C,D).

**Figure 6:**
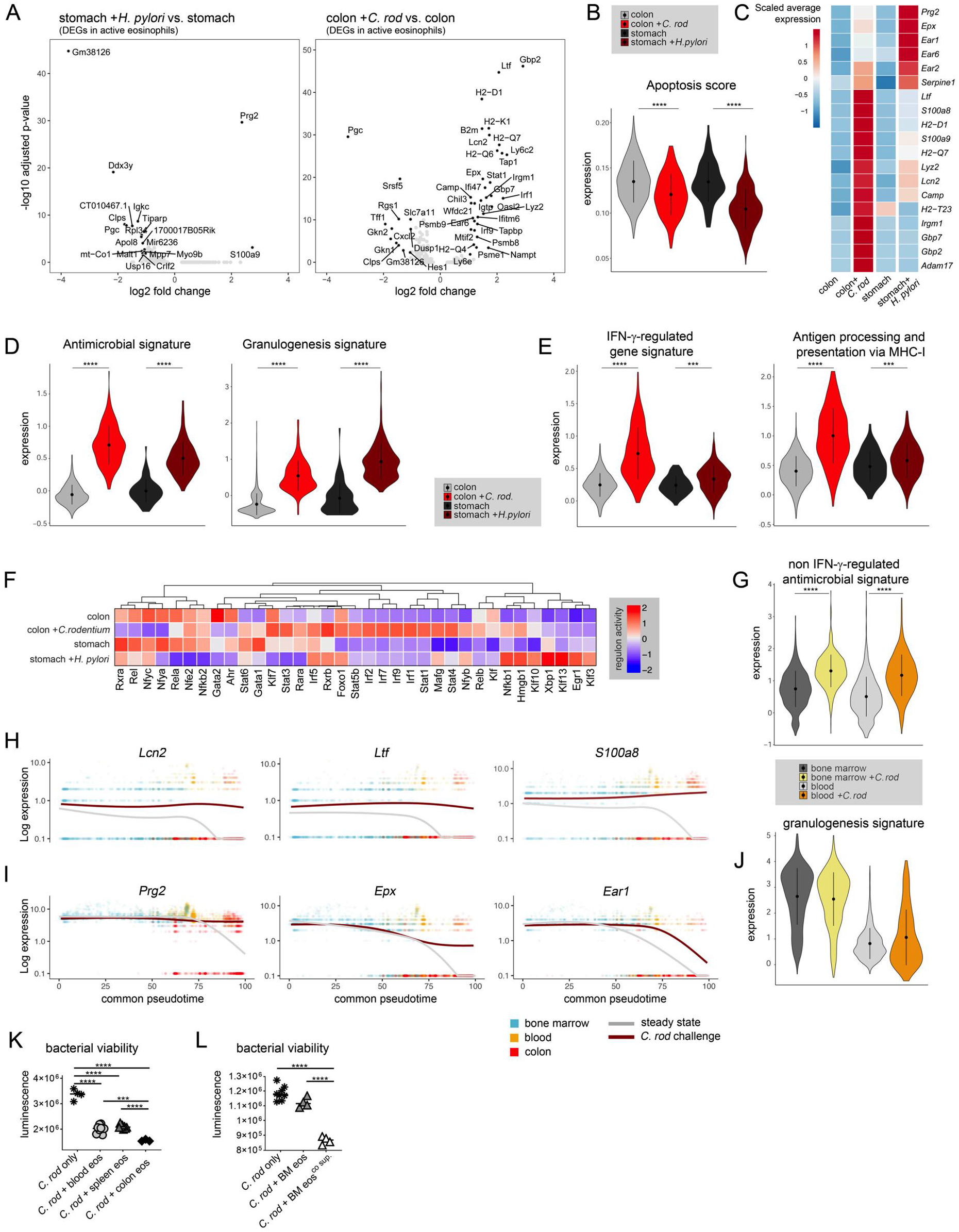
Challenge infection promotes antimicrobial gene programs in active eosinophils. **A)** Significant DEGs within the active subset (average log fold-change > 0.5, adjusted P < 0.05), at steady state and during bacterial infection. **B)** Apoptosis score in active eosinophils. **C,D)** Antimicrobial and granulogenesis genes (C) and signatures (D) in active eosinophils. **E)** Expression of IFN-γ-regulated and MHC-I-restricted antigen processing and presentation signature genes in active eosinophils. **F)** Regulon activity in active eosinophils across conditions. **G)** Non-IFN-γ regulated antimicrobial signature gene expression in BM and blood, at steady state and during ***C.** rod* infection. **H-I)** Gene expression over aligned pseudotime at steady state (gray) and during ***C.** rod* infection (dark red). Dots indicate single cells, colored by organ. **J)** Granulogenesis signature in BM and blood at steady state and during ***C.** rod* infection. **K-L)** ***C.** rod* (ICC180) viability upon 60 min. exposure to blood, splenic and colonic eosinophils (K) or upon exposure to BM-derived eosinophils, conditioned with colonic supernatant or left untreated (L). Technical replicates and medians are shown. Eosinophils were pooled from n = 3 *Il5*-tg (K) or B6J (L) mice. ***P < 0.001; ****P < 0.0001 (One-way ANOVA). B, D, E, G, J) Genes used for scores and signatures are listed in Table S2. Data represent the mean ± SD. ***P < 0.01; ****P < 2.2×10^−16^ (two-sided Wilcoxon test). See also Figure S6.

In accordance with the robust type I immune response triggered by *C. rod*, active eosinophils exhibited marked IFN-γ-regulated gene expression (Figures 6E and S6C), while this signature was weaker in the early stages (4 weeks) of chronic *H. pylori* infection. The significant enrichment of terms related to MHC-I-restricted antigen processing and presentation further indicates crosstalk with CD8^+^ T cells (Figures 6E and S6B). Indeed, co-culture of eosinophils with OT-I CD8^+^ T cells, but not OT-II CD4^+^ T cells, resulted in robust T cell proliferation in an antigen-dependent manner, suggesting that BM-eos can present antigen via MHC-I/TCR interactions. This was however independent of their prior conditioning into the active subset with colon supernatant (Figure S6D).

We further examined the impact of infection on the TF network expression in active eosinophils by SCENIC (Figure 6F). TFs governing transcriptional responses during *C. rod* infection included multiple STATs (Stat1, 3, 4, 5b, 6), IRFs (Irf1, 2, 5, 7, 9) and TFs of the NF-κB family (Nfkb1, Nfkb2, Relb), indicating signaling inputs from a broad range of cytokines. High IRF1 and Stat1 regulon activity further supported the emergence of an IFN-γ-mediated gene signature, which was predicted to directly upregulate granule and antimicrobial peptide-related gene programs.

Bacterial challenge further affected the expression of multiple transcripts in the BM and blood (Table S1), including IFN-γ-independent antimicrobial genes (Figures 6G and S6E). To identify differentially expressed genes (DEG) throughout eosinophil development, we computed BM-blood-colon trajectories during steady state and bacterial challenge, and aligned them to a common pseudotime axis by dynamic time warping (McFaline-Figueroa et al., 2019). Among the 149 DEGs in response to *C. rod* infection, antimicrobial peptides such as *Lcn2*, *Ltf*, *S100a8*, *S100a9*, *Camp* and *Lyz2* were already upregulated in the BM, with high expression maintained at every stage of maturation (Figures 6H and S6F). In contrast, eosinophil cytotoxic granule protein expression in the BM remained unchanged during infection, but was consecutively maintained in the blood and colon (Figures 6I,J and S6G), and was independent of stemness and proliferation programs (Figure S6H, I). We further defined several IFN-γ-dependent genes, which upregulation was limited to the colon. These included the antimicrobial peptide *Gbp7* and the MHC-I molecule component *B2m* (Figure S6J). Of note, *Ifngr1* expression was downregulated in response to infection (Figure S6J), potentially as a feed-back mechanism to fine-tune eosinophil activation (Crisler et al., 2019). In addition, IFN-γ treatment upregulated *Cd274* expression directly in BM-eos, particularly in conjunction with *C*. *rod* infection (Figure S6K).

Our data indicate that active eosinophils increase their antimicrobial and cytotoxic protein arsenal in response to bacterial challenge. To probe eosinophil bactericidal activity *in vitro*, purified blood, spleen and colon eosinophils were incubated with a bioluminescent *C. rod* strain, which allows the assessment of bacterial viability over time through quantification of the bioluminescent signal (Figure S6L). As expected, all three eosinophil populations were able to limit *C. rod* growth, but colonic eosinophils exhibited significantly greater bactericidal activity than the other two populations (Figure 6K). In addition, BM-eos required prior conditioning with colonic supernatant for optimal bacterial killing (Figure 6L), an effect that was not recapitulated by conditioning with IFN-γ, TNF-α or IL-17 alone (Figure S6M).

### Eosinophils protect mice from experimental colitis and active eosinophils accumulate in IBD patients

We next sought to confirm our findings in an additional model of colitis, induced by dextran sulfate sodium (DSS). We retrieved eosinophil transcriptomes from an independent BD Rhapsody dataset profiling colonic CD45+ cells of DSS-treated B6J mice (Schwazfischer et al., submitted). After integration into our dataset, 99% of all colonic eosinophils mapped to the active cluster (Figure 7A) and expressed the characteristic transcripts *Cd9*, *Cd80* and *Cd274* (Figure S7A). In accordance with our observations during bacterial challenge, active eosinophils upregulated genes associated with MHC-I-restricted antigen processing and presentation (Figure 7B), as well as IFN-γ-related transcripts and antimicrobial peptides (Figure 7C). In independent experiments, DSS treatment induced a striking increase in the frequency of PD-L1^+^ CD80^+^ active eosinophils (Figure 7D), while eosinophil deficiency (PHIL mice) resulted in increased colitis severity (Figure 7 E, F), as reported previously (Masterson et al., 2015). PHIL mice also exhibited stronger Th17 responses relative to their WT littermates, as well as increased TNF-α production by CD4^+^ T cells (Figure S7B), indicating an immune regulatory role of active eosinophils in DSS-induced colitis. Indeed, co-culture of naïve BM-eos with CD4^+^ T cells resulted in enhanced T cell proliferation following anti-CD3/CD28-mediated stimulation, while the prior eosinophil conditioning with colonic supernatant induced significantly lower T cell proliferation and was further reduced by supplementing the conditioning medium with IFN-γ (Figure 7G). By applying the cell-cell interaction prediction tool CellPhoneDB (Efremova et al., 2020) to eosinophils, CD4^+^ and CD8^+^ T cells in the DSS dataset, we further identified numerous potentially interacting ligand-receptors pairs (Figure S7C), indicative of sustained crosstalk between these cell types during inflammation.

**Figure 7:**
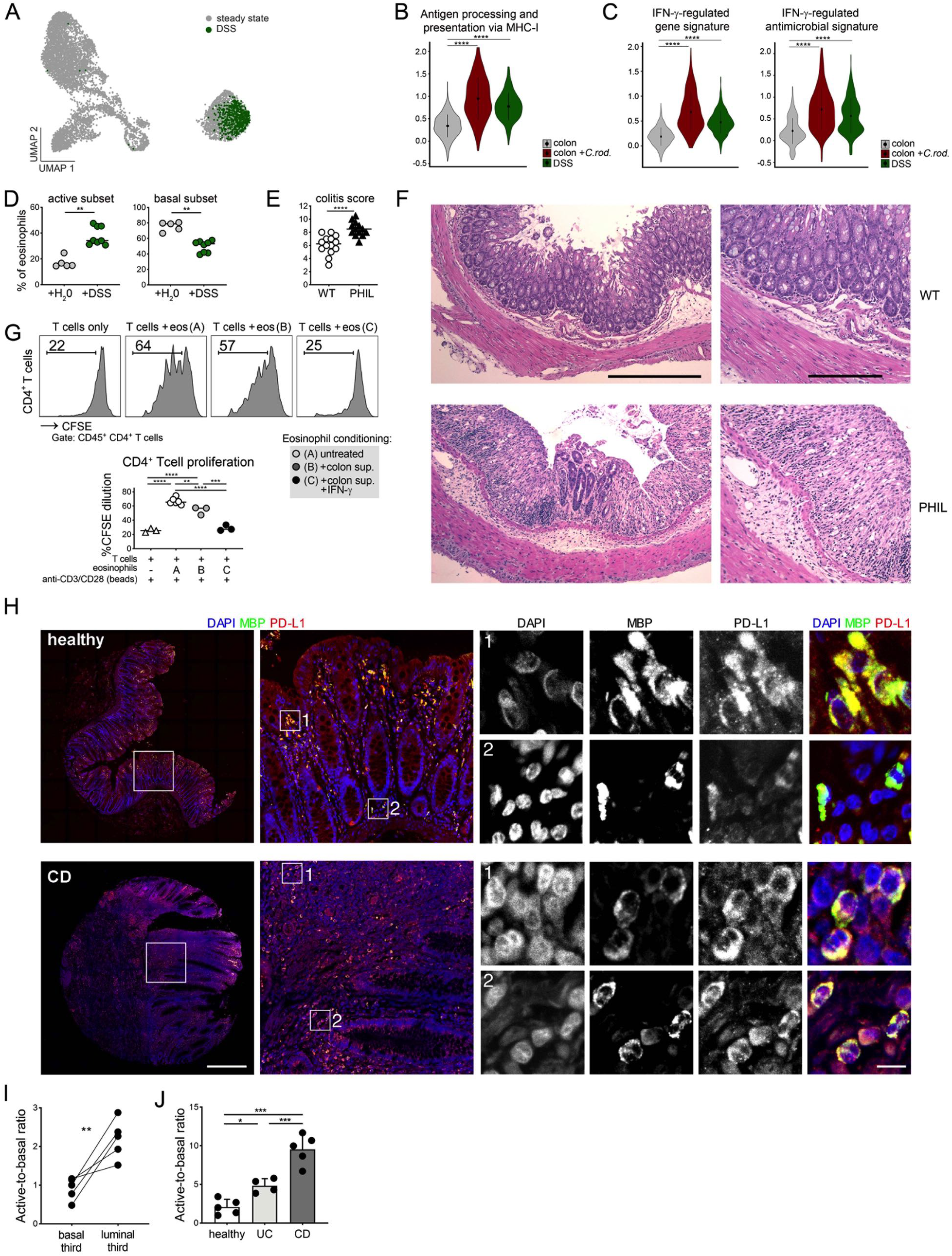
Eosinophils protect mice from experimental colitis and active eosinophils accumulate in IBD patients. **A)** Data integration of colonic eosinophils isolated from DSS-treated B6J mice (green dots, n = 3). Steady state dataset (gray dots) is used as a reference. **B-C)** Expression of MHC-I-restricted antigen processing and presentation (B), IFN-γ-regulated genes and antimicrobial signature genes (C) in active eosinophils. Genes used for scores and signatures are listed in Table S2. Data represent the mean ± SD. ****P < 2.2×10^−16^ (two-sided Wilcoxon test). **D)** Frequencies of active and basal eosinophils in the colon of DSS-treated vs naïve B6J mice (n = 5-8); medians are shown. **E-F)** Colitis score in B6J and PHIL mice assessed by histopathological examination (E) and representative H&E-stained colonic sections (F). (n = 13-14); data are pooled from two independent experiments. Medians are shown. Scale bars, 100 *μ*m. **G)** Anti-CD3/CD28-activated, CFSE-labelled naïve CD4^+^ T cells were co-cultured with B6J BM-eos that were conditioned with colon supernatants ± IFN-γ or were left untreated. Representative flow cytometry plots (top) and quantification (bottom) of T cell proliferation assessed by CFSE dilution. BM-eos were generated from n = 2, B6 mice. Medians are shown. Data representative of three independent experiments. **H)** MBP and PD-L1 immunofluorescent staining of human tissue microarrays. Representative cores from healthy individual and CD patient are shown. Scale bars, 500 *μ*m (core overview) and 10 *μ*m (high magnification insets). **I)** Active-to-basal ratio in luminal (upper) third vs. basal (lower) third colonic crypts quantified in healthy human colon cores (n = 5). ** P < 0.01 (unpaired t-test) **J)** Active-to-basal eosinophil ratio in healthy individuals (n = 5), CD (n = 5) and UC patients (n = 4). D,E) **P < 0.01; ****P < 0.0001 (Mann-Whitney test). G,J) *P < 0.1; **P < 0.01 ***P < 0.001; ****P < 0.0001 (One-way ANOVA). See also Figure S7.

Finally, we investigated the existence of a human equivalent of murine active GI eosinophils. As eosinophils are not included in the human intestinal scRNA-seq datasets published so far (Smillie et al., 2019), we stained healthy and IBD patient tissue microarrays for eosinophil major basic protein (MBP). Both MBP^+^ PD-L1^+^ (active) and MBP^+^ PD-L1^−^ (basal) eosinophil subsets were found in the human LP (Figure 7H), with active eosinophils located predominantly in the luminal section of the mucosa (Figure 7I), similar to their murine equivalents. Interestingly, the active-to-basal eosinophil ratio was 5-fold enriched in CD specimens, but only 2-fold enriched in ulcerative colitis (UC) patients relative to healthy controls (Figure 7J), suggesting active eosinophils also accumulate in IBD patients, especially in CD. The transcriptomic profile of MBP^+^ PD-L1^+^ (active) eosinophils, as well as their role in the pathogenesis of IBD warrants further investigation.

## Discussion

The extended survival and high numbers of eosinophils present in the GI tract at steady state, and their accumulation during infection and inflammation, suggest eosinophils play cardinal roles in intestinal homeostatic and inflammatory processes. Here, we investigated the existence and ontogenetic relationship of eosinophil subsets to define their functional contribution to intestinal health and diseases. We identified five consecutive developmental and maturation stages of the eosinophil lineage: (i) progenitors in the BM, the only proliferative stage, (ii) immature eosinophils, characterized by granule protein production and present in the BM and blood, (iii) circulating eosinophils that migrate to target tissues, (iv) basal eosinophils, a tissue-resident population endowed with morphogenetic properties and (v) GI active eosinophils, characterized by PD-L1 and CD80 expression, bactericidal activity and tissue-protective function during inflammation.

Our findings suggests that eosinophil subpopulations emerge from consecutive maturation steps and adaptation to distinct niches. This observation corroborates proposals that the considerable morphological and phenotypic variability of tissue eosinophils reflects their adaptation to different microenvironments (Abdala-Valencia *et al.*, 2018; Masterson *et al.*, 2021). In the GI tract, active, but not basal eosinophils exhibited strong transcriptional and phenotypical heterogeneity across compartments, suggesting this subset readily adapts to local microbial, tissue and cytokine cues. The residence of active eosinophils in the luminal third of the intestinal mucosa coincides with the activation of HiF1-α, which may allow their adaptation to the oxygen gradient that decreases steeply from the intestinal submucosa to the anaerobic lumen (Zheng et al., 2015). The presence of this subset close to the intestinal epithelium is consistent with a CD11chi eosinophil population reported to localized to the intraepithelial fraction of the SI (Xenakis *et al.*, 2018). Thus, our study expands the concept of myeloid cell plasticity and tissue adaptation to the eosinophil lineage, a notion so far only explored thoroughly for macrophages (Lavin et al., 2014) and neutrophils (Ballesteros et al., 2020).

Our transcriptional analysis implicates both basal and active eosinophils in tissue morphogenesis, homeostasis and immune regulation at steady state. Basal eosinophils express *Tgfb1,* which is cleaved into its active form by MMP9 and THBS1 (Crawford et al., 1998; Kobayashi et al., 2014), two enzymes encoded by the active and basal subsets, respectively. GI eosinophils may thus be a prominent source of active TGF-β, promoting fibroblast proliferation, wound healing and regulatory T cell differentiation (Levi-Schaffer et al., 1999; Phipps et al., 2002) (Chu *et al.*, 2014). The upregulation of multiple enzyme, cytokine and chemokine transcripts in active eosinophils further indicates a marked increase in biosynthetic activity upon differentiation of this subset. Although eosinophil cytokines are predominantly found preformed and stored within their granules and secretory vesicles (Melo et al., 2013), *de novo* production of these factors in the active cluster suggests they are strictly required to fulfill tissue-specific activities. Among the signature genes of this cluster, *Vegfa, Il1b* and *Il1rn* have been linked to eosinophil-mediated angiogenesis (Hoshino et al., 2001), support of intestinal homeostasis and IgA production (Jung *et al.*, 2015) and local Th17 response downmodulation (Sugawara *et al.*, 2016), respectively. Our study further highlights the prevalent expression of T cell co-stimulatory molecules by basal (*Icosl*) and, more predominantly, by active eosinophils (*Cd9*, *Cd80*, *Cd274*) and identifies CD80 and PD-L1 as *bona fide* markers of both the human and mouse active eosinophil subset. We indeed show that eosinophils enhance T cell proliferation in their basal state, but downmodulate this activity following intestinal conditioning, particularly upon IFN-γ exposure. Active eosinophils might thus control the extent of CD4^+^ T cell responses, potentially via co-stimulatory molecule expression, to limit tissue damage during Th1 inflammation. These observations are consistent with our previous report indicating a functional role for PD-L1 and IFN-γ signaling in eosinophil-mediated T cell suppression (Arnold *et al.*, 2018) and corroborate studies supporting the regulation of T cell activities by GI eosinophils (Arnold et al., 2020; Sugawara *et al.*, 2016). Further functional characterization of the eosinophil co-stimulatory markers identified in this study (CD80, CD9, ICOSL) is warranted.

Among the signals driving active eosinophil differentiation, we identified microbially-derived cues, NF-κB signaling and IFN-γ signaling during infection, as critical drivers of their maturation *in situ*. Active eosinophil differentiation required microbial presence but was independent of major bacterial recognition pathways (MyD88, TLR2/4), suggesting that microbiota-derived signals act indirectly on eosinophils via the epithelium or other immune cells. Direct contacts between eosinophils and commensals are indeed unlikely to occur under homeostatic conditions, given the strict compartmentalization of the microbiota within the intestinal lumen. While we did not identify a single factor sufficient to drive active eosinophil differentiation, this process may require synergistic signals. Additional microbiota-regulated factors known to signal at least partially via the NF-κB pathway may include the epithelial-derived cytokine IL-25 (Buonomo et al., 2016), hypoxia (Culver et al., 2010) and Ahr ligands (Ovrevik et al., 2014), but require further investigation.

An intriguing new finding of our study is the striking compositional shift toward the active cluster at sites of infection and inflammation, which coincides with the acquisition of antimicrobial functions. The absence of new eosinophil cluster(s) arising during bacterial infection and colitis, as observed in three independent models, suggests that eosinophils recruited in this context do not differ substantially from their steady state counterparts, but rather acquire new functional properties, largely driven by local interferon and cytokine signaling networks (Irf1/2/7/9; Stat1/3-6; NFkb1/2, Rela/b). This finding contrasts with a previous report indicating that lung-resident eosinophils differ morphologically, phenotypically and functionally from those accumulating during allergen challenge (Mesnil *et al.*, 2016). Interestingly, the upregulation of antimicrobial peptides and cytotoxic granule protein genes is already detectable during eosinophil maturation in the BM and blood. The sustained expression of antimicrobial gene programs throughout eosinophil development, together with the bypassing of certain maturity stages, such as the basal state, may thus ensure the rapid deployment of “primed” eosinophils at sites of bacterial invasion. The increased survival of active eosinophils during infection, as shown by their decreased expression of apoptosis-related genes, is consistent with our previous report showing that eosinophils undergo “non-lethal” degranulation and EET release in response to *C. rod*, thus promoting bacterial clearance and host protection (Arnold *et al.*, 2018). The presence of a PD-L1^+^ eosinophil subset in the normal human mucosa, and its accumulation during IBD, suggests the existence of a human counterpart of the active eosinophil subset that may share its functional properties.

Overall, our study further elucidates the ontogeny of intestinal eosinophils and reveals the marked transcriptomic changes that occur during eosinophil maturation across anatomical compartments. The gene programs upregulated by eosinophils in the healthy and inflamed GI tract further highlight the major contribution of these cells to intestinal homeostasis, immune regulation and host defense. Our findings provide novel insights into the biology of this elusive cell type and may lay a framework for the functional characterization of eosinophil subsets in GI diseases.

### Limitations of the study

A potential limitation of our study is that we did not profile eosinophils of other mucosal tissues to confirm the GI specificity of the active subset. In addition, we have not investigated whether the predominant expression of regulatory and bactericidal gene programs of active eosinophils would also emerge in the context of Th2 inflammation. Consequently, we cannot rule out a potential correspondence between the GI active subset and the recently described inflammatory eosinophils recruited to the lungs during allergen challenge (Mesnil *et al.*, 2016), despite limited phenotypical similarities. Further studies are required to assess whether active eosinophils equivalents exist in other tissues and whether their transcriptomic profile is conserved during Th2 inflammation.

## Supporting information

Table S2. List of genes used to define scores and signatures, with respective references.

Table S3. List of antibodies (antigen, clone, fluorochrome, dilution and manufacturer) used for high dimensional spectral flow cytometric analysis.

## Acknowledgements

We thank K. Stirm, S. Baghai Sain and A. Lafzi, and A. Müller for technical support, discussion and insightful ideas. This study was supported by an SNSF Eccellenza Professorial Fellowship from the Swiss National Science Foundation (SNF) to I.C.A. (PCEFP3_187021) and A. E. M (PCEFP3_181249); A.G. is supported by fellowship 2450 from the Hartmann-Müller Foundation; I.G.P is supported by fellowship 20C197 from the Novartis Foundation for Medical-Biological Research. T.V. is supported by Czech Science Foundation grant 21-26025S; T.V and N.G.N are fellows of the University of Zürich Research Priority Program (URPP) “Translational Cancer Research”. This paper was typeset with the bioRxiv word template by @Chrelli: www.github.com/chrelli/bioRxiv-word-template

## Author contributions

A.G., C.B., I.G.P., N.G.N., D.C., K.H. and T.V. performed experiments and analyses. C.B., N.G.N. and K.H. performed bioinformatics analyses. I.C.A., A.E.M, B.B., and K.B. designed and supervised experiments. I.C.A., C.B. and A.G. wrote the manuscript.

## Materials and Methods

### Mice

All experiments were performed on 6-12 week-old male and female mice. The following mice were used: C57BL/6J (B6J, stock no. 000664) obtained from The Jackson Laboratory; OT-1 (stock no. 003831), OT-II (stock no. 004194), MyD88^-/-^ (Adachi et al., 1998), Tlr2^-/-^ (stock no. 004650), TNFR1^-/-^ (stock no. 002818), CD45.1 (stock no.002014), IFN-γR^-/-^ (stock no.003288) and Tlr4 ^-/-^ (Hoshino et al., 1999) were obtained from a local live mouse repository. *Il5*–transgenic mice have been previously described (Dent et al., 1990). Eosinophil-deficient mice (PHIL) were obtained from J.J. Lee (Mayo Clinic, Phoenix, AZ) and described previously (Lee et al., 2004). Chow and water were available *ad libitum*, unless specified. All mice were in the B6J background and maintained on a 12h light / 12h darkness schedule. Mice were housed and bred under specific pathogen-free conditions in accredited animal facilities. At the experimental endpoint, mice were sacrificed by raising CO2 concentrations. All experimental procedures were performed in accordance with Swiss Guidelines and approved by the Cantonal Veterinary Office Zürich, Switzerland and/or in accordance with the European Communities Council Directive (86/609/EEC), Czech national guidelines, institutional guidelines of the Institute of Molecular Genetics and approved by the Animal Care Committee.

### Animal treatments

#### Antibody neutralization

7-8 week-old B6J females were injected intraperitoneally (i.p.) with a single dose of 0.5mg anti-IL-5 (BE0198 BioXCell, TREK5) or anti-GM-CSF (BE0259 BioXCell, MP1-22E9) antibodies, or with anti-horseradish peroxidase isotype control (BE0088 BioXCell, HRPN), 3 days before the study endpoint.

#### Intestinal commensal depletion by antibiotic treatment

7-8 week-old B6J females were treated for 10 consecutive days with ampicillin (1 g/L; A0166 Sigma), vancomycin (500 mg/L; A1839,0001 Applichem), neomycin sulfate (1 g/L; 4801 Applichem), and metronidazole (1 g/L; H60258 Alfa Aesar) in autoclaved drinking water, as previously described (Diehl et al., 2013). Water bottles were monitored and refilled twice per week.

#### Adoptive transfer

10^6^ cells magnetically-selected splenic eosinophils of 6-12 week-old *Il5*-tg females and males were injected intravenously in 100ul PBS into CD45.1 recipients (8-12 week-old female and male mice). Organs were harvested 42 hours after injection.

#### DSS-induced colitis

6-12 week-old females and males were treated with 2.5% Dextran Sulfate Sodium (DSS) (wt/vol; 9011-18-1, MP Biomedicals) dissolved in autoclaved drinking water for 5 days, followed by 3 days of regular water, before organ harvesting. Water bottles were monitored and refilled twice per week.

#### Bacterial challenge infection, H. pylori

6-12 week-old *Il5*-tg or B6J females and males were infected orally with *H. pylori* strain PMSS1 (107 colony-forming units, CFU) and analysed 4 weeks post-infection. The PMSS1 strain, a clinical isolate of a duodenal ulcer patient, was grown on horse blood agar plates followed by liquid culture, as previously described (Arnold et al., 2011). Cultures were routinely assessed by light microscopy for contamination, morphology and motility. *C. rodentium:* 6-12 week-old *Il5*-tg or B6J females and males were infected orally with the nalidixic acid-resistant *C. rod* strain ICC169 (ATCC 51549, 108 CFU) and analyzed 13 days post-infection. Bioluminescent *C. rodentium* strains ICC180 (ICC169 derivative, nalidixic acid- and kanamycin-resistant) was a kind gift of Gad M. Frankel, Imperial College London, UK and was previously described (Wiles et al., 2006). Both strains were grown on agar plates (1.5%; A0927 Applichem), followed by single-colony picking and overnight culture in antibiotic-supplemented Luria broth (nalidixic acid, 50 μg/ml; N4382 Sigma and/or kanamycin, 50 μg/ml; 420311 Sigma).

### Preparation of single-cell suspensions from tissues

#### Gastrointestinal tissues

stomach, colon and small intestine (SI) were harvested, cleaned of faecal matter and cut longitudinally. Organs were washed in PSB and cut into pieces (1-2cm) and Peyer’s patches were removed from the SI. Pieces were washed twice in a shaking incubator with wash buffer (2% BSA, 100 U/mL penicillin/streptomycin, 5 mM EDTA in HBSS, 25 minutes, 37 °C). Tissues were then rinsed in cold PBS and digested for 50 minutes at 37°C in complete medium (10% FBS, 100 U/ml, penicillin/streptomycin (P0781 Sigma) in RPMI-1640) containing 15 mM Hepes (H0887 Sigma), 0.05 mg/ml DNase I (10104159001 Roche) and an equal amount of 250 U/mL type IV (C5138 Sigma) and type VIII collagenase (C2139 Sigma) (for colon and SI), or 500 U/mL type IV collagenase (for stomach). Cells were passed through a 70μm cell strainer, centrifuged for 8 minutes and layered onto a 40/80% Percoll (17089101 Cytiva) gradient (18 minutes, 2100 g, 20°C, no brake). The interphase was collected and washed in PBS.

#### Lung

lungs were perfused with PBS, harvested and cut into pieces before digestion in complete medium supplemented with 500 U/mL type IV collagenase (Sigma) and 0.05 mg/ml DNase I (Roche) for 50 minutes at 37°C. Lungs were then passed through a 70μm cell strainer and mesh with syringe plungers.

#### Blood

blood was sampled by post-mortem cardiac puncture in 2% BSA 5mM EDTA PBS. The suspension was layered over Histopaque 1119 (density of 1.119 g/mL; 11191 Sigma-Aldrich) and centrifuged at 800g for 20 minutes. The interphase was washed in PBS and red blood cells were lysed in ice-cold distilled water for 30 seconds.

#### Bone marrow (BM)

femur and tibia were flushed through a 23-gauge needle and collected in complete RPMI medium. Suspensions were filtered through a 40μm cell strainer and red blood cells were lysed in ice-cold distilled water for 30 seconds.

#### Spleen and lymph nodes

spleen and lymph nodes were harvested, meshed through a 40μm cell strainer using a syringe plunger, and red blood cells were lysed in ice-cold distilled water for 30 seconds.

Unless specified, all centrifugation steps were performed at 500 g for 8 minutes at 10°C.

### Eosinophil enrichment by magnetic selection

Eosinophils of 6-12 week-old *Il5*-tg females and males were positively-enriched using PE anti-mouse Siglec F antibody (562068 BD Biosciences; E50-2440) and anti-PE microbeads (130-042-401 Miltenyi Biotech), according to the manufacturer’s instructions.

### Single-cell RNA sequencing

Whole transcriptome analyses were performed on magnetically-enriched Siglec F^+^ eosinophils (blood, spleen, stomach, colon and SI) and total bone marrow cells using the BD Rhapsody Single-Cell Analysis System (BD, Biosciences). Cells were pooled from 3-4 7-8 week-old *Il5*-tg females and males per sample. Tissue processing and enrichment procedures are described above. Each preparation was assessed by flow-cytometry to determine eosinophil viability and was subjected to morphological examination upon cytospin and staining. Eosinophils were labelled with sample tags (633793 BD Mouse Single-Cell Multiplexing Kit) according to the the manufacturer’s protocol. Briefly, for each condition, 106 cells were resuspended in staining buffer (1% BSA, 1% EDTA in PBS) and incubated with the respective Sample Tag for 20 minutes at room temperature. Cells were then transferred to a 5 mL polystyrene tube, washed two times with 2 mL staining buffer and centrifuged at 400g for 5 minutes. Samples were resuspended in 1 mL staining buffer for counting. 10’000 or 20’000 cells from up to 4 barcoded samples were pooled for a total of 60’000 cells, the mixture was centrifuged at 400g for 5 minutes. The pellet was resuspended in 650 BD Sample Buffer supplemented with 1:1000 SUPERase in (20U/*μ*L; AM2694 ThermoFisher) and NxGen Rnase Inhibitor (40U/*μ*L; 30281-2 Lucigen). BD Rhapsody cartridges were super-loaded with 60’000 cells each. Single cells were isolated with the BD Rhapsody Express Single-Cell Analysis System according to the manufacturer’s recommendations (BD Biosciences). cDNA libraries were prepared using the BD Rhapsody Whole Transcriptome Analysis Amplification Kit (633801 BD Biosciences) following the BD Rhapsody System mRNA Whole Transcriptome Analysis (WTA) and Sample Tag Library Preparation Protocol (BD Biosciences). The final libraries were quantified using a Qubit Fluorometer with the Qubit dsDNA HS Kit (Q32851 ThermoFisher). Library size-distribution was measured with the Agilent high sensitivity D5000 assay on a TapeStation 4200 system (5067-5592 Agilent technologies). Sequencing was performed in paired-end mode (2*75 cycles) on a NovaSeq 6000 with NovaSeq 6000 SP Reagent Kit chemistry.

### Flow cytometry, cell sorting and counting

For surface staining, cells were stained in PBS at 4°C for 30 minutes with the fixable viability dye eFluor 780 (1:1000, 65-0865-14 eBioscience) and a combination of the following antibodies (1:200, all from BioLegend unless stated otherwise): anti-mouse CD45 BV650 (30-F11, 103151), CD11b BV510 (M1/70, 101263), MHC-II AF700 (M5/114.15.2, 107622), Ly6G Percp-Cy5.5 (1A8, 127616), CD4 PerCP (RM4-5, 100538), TCRβ PE-Cy7 (H57-597, 109222), CD80 BV605 (16-10A1, 104729), PD-L1 PE-Cy7 (10F.9G2, 124314), CD31 PE (390, 102408), CD45.2 BV785 (104, 109839), CD9 PE (MZ3, 124805), CD54 BV711 (YN1/1.7.4, 116143), CD95 PE-Cy7(SA367H8, 152607), SiglecE PE (M1304A01, 677104), Sca-1 AF488 (D7, 108116), CD11c APC-Cy7 (N418, 117323), Clec12a PE (5D3, 143404), CD49d FITC (R1-2, 103605), CD16/32 FITC (S17012B, 101305), CD3e Percp-Cy5.5 (145-2C11, 100328), CD8a APC (53-6.7, 100712), Siglec F BV421 (E50-2440, 552126 BD Biosciences), CD275 (HK5.3, 50598582 eBioscience). For T cell intracellular cytokine staining, cells were incubated for 3.5 h in complete IMDM medium containing 0.1 μM phorbol 12-myristate 13-acetate (P-8139 Sigma) and 1 μM ionomycin (I-0634 Sigma) with 1:1000 Brefeldin A (00-4506-51 eBioscience) and GolgiStop solutions (51-2092KZ BD Biosciences) in a humidified incubator with 5% CO2, 37°C. Following surface staining, cells were fixed and permeabilized with the Cytofix/Cytoperm Fixation/Permeabilization Solution kit (512090KZ BD Biosciences) according to the manufacturer’s instructions. Cells were then stained for 50 minutes with anti-mouse IL-17A APC (TC11-18H10.1, 506916), IFN-γ BV421 (XMG1.2, 505830) and TNF-ɑ FITC (MP6-XT22, 506 304) all from Biolegend. Fc block (anti-CD16/CD32, 101302 Affymetrix) was included to minimize nonspecific antibody binding. Total leukocyte counts were determined by adding countBright Absolute Counting Beads (C36950 Life Technologies) to each sample before analysis. Samples were acquired in a LSRII Fortessa or FACS AriaIII 5L (BD Biosciences). For high-dimensional spectral flow-cytometry analysis, cells were acquired on Cytek Aurora 5L (Cytek Biosciences) following 50 minutes staining at 4°C with the antibodies described in Table S3. BD FACSDiva Software (BD Biosciences) was used for data acquisition and cell sorting.

### Isolation and culture of mouse BM-derived eosinophils

To generate murine BM-derived eosinophils (BM eos), BM cell suspensions were seeded at a density of 10^6^ cells/mL in RPMI-1640 medium supplemented with 20% heat inactivated FBS, 25 mM Hepes (H0887 Sigma), 100 U ml−1 penicillin/streptomycin (P0781 Sigma), 2 mM glutamine (25030-024 Gibco), 1xNEAA (11140-035 Gibco), and 1 mM sodium pyruvate (11360070 Gibco). Cells were cultured in a humidified incubator with 5% CO_2_, 37°C, and were supplemented with 100 ng/ml mouse SCF (250-03 PeproTech) and 100 ng/ml mouse FLT3-Ligand (250-31L PeproTech) from day 0 to day 4, followed by differentiation with 10ng/mL murine IL-5 (215-15 PreproTech) until day 13, as described (Dyer et al., 2008). Half of the medium was replaced and the cell concentration was adjusted to 10^6^ cells/mL every other day. On day 8, cells were collected and moved to new flasks to remove adherent contaminating cells. On day 13, the nonadherent cells were collected and washed with PBS. Eosinophil purity was assessed by flow cytometry (>95%).

### *In vitro* conditioning with supernatant of cultured colonic explants

Supernatant of cultured colonic explants (colon supernatant) was prepared by culturing mid colon sections (~0.3 cm) from 6-12 week-old B6J females and males in 300*μ*L of complete RPMI medium in a humidified incubator with 5% CO_2_, 24 hours at 37°C. Blood and splenic eosinophils were magnetically isolated from *Il5*-tg mice or differentiated from the BM of B6J mice and were kept in complete RPMI medium with recombinant murine IL-5 (10ng/mL, 215-15 PreproTech). Cells were seeded in round-bottom 96-well plates at a density of 2×10^5^ cells / well (100*μ*L) and conditioned for 12 hours at 37°C with cell-free supernatant (1:10).

### *C. rodentium* ICC180 viability assay

Flow-cytometry-purified BM-eos (B6J) and magnetically-enriched colonic, splenic and blood eosinophils (*Il5*-tg) from 6-12 week-old females and males were conditioned overnight and/or treated with recombinant mouse cytokines, as indicated (IFN-γ, 10ng/mL, 315-05; TNF-a 20ng/mL, 315-01; IL-17A 10ng/mL, 210-17; all from PeproTech). Eosinophils were washed with PBS and transferred to a solid white flat bottom 96-well plate (Corning) in antibiotic-free RPMI-1640 medium supplemented with 10% FBS and murine IL-5 (10 ng/mL, PeproTech). 10^8^ bioluminescent *C. rod* bacteria (at exponential phase, 1–1.5 OD600) were added to each well and luminescence was measured after 2 minutes and subsequently every 10 minutes for the following 2 hours on an Infinite 200 PRO plate reader (TECAN).

### T cell proliferation assay

Flow-cytometry-purified BM-eos (B6J) were conditioned overnight with colon supernatant and treated with recombinant mouse IFN-γ (10ng/ml, 315-05 PeproTech), as indicated. Naïve CD4^+^ T cells were isolated from the lymph nodes, enriched with the MojoSort Mouse CD4 Naïve T Cell Isolation Kit (480040 BioLegend) and were purified by flow-cytometry. T cells were labelled with the CellTrace CFSE Cell Proliferation Kit (C34554 ThermoFisher) following manufacturer’s instructions. T cells were then activated by CD3/CD28 T-activator Dynabeads (11131D Gibco) and co-cultured with eosinophils at a 1:1 ratio (2×10^5^ total) for 4 days at 37°C in complete RPMI medium supplemented with 10 ng/mL recombinant mouse IL-5 (PeproTech) and 20 ng/mL IL-2 (402-ML R&D). CFSE dilution was assessed by flow cytometry.

### Antigen presentation assay

Flow-cytometry-purified BM-eos (B6J) were conditioned overnight with colon supernatant, where indicated. Cells were washed in PBS and loaded with 300 ng/mL of ovalbumin (OVA) residues 257-264 (S7951 Sigma) or 323-339 (O1641 Sigma) for 6 hours in complete RPMI medium supplemented with 10 ng/mL recombinant IL-5 (PeproTech). T cells were sorted by flow-cytometry and labelled with CellTrace CFSE Cell Proliferation Kit (C34554 ThermoFisher) following manufacturer’s instructions. OT-I CD8^+^, OT-II CD4^+^, and CD8^+^ / CD4^+^ WT T cells were obtained from the lymph nodes of 8-12 week-old female and male OT-I, OT-II and B6J mice, respectively. T cells were co-cultured with eosinophils at a 1:1 ratio (2×10^5^ total) for 4 days at 37°C in complete RPMI medium supplemented with 10 ng/mL recombinant mouse IL-5 (PeproTech) and 20 ng/mL IL-2 (402-ML R&D). CFSE dilution was assessed by flow cytometry.

### Quantitative RT-PCR

The RNA of tissue-derived or cultured BM-eos was isolated using Directzol RNA MicroPrep kit (R2062 Zymo Research) according to the manufacturer’s instructions, including a DNase I digestion step. Complementary DNA synthesis was performed using Superscript III reverse transcription (18080-044 QIAGEN). Gene expression was measured on a CFX384 Touch Real-Time PCR system (BioRad, Second Derivative Maximum method analysis with high confidence algorithm) by TaqMan Gene Expression Assays (4331182 Applied Biosystems by ThermoFisher Scientific): *Cxcl2* (Mm00436450_m1), *Hprt* (Mm03024075_m1), *Gapdh (Mm99999915_g1), Cd274* (Mm03048248_m1)*, Cd80 (Mm00711660_m1), Ahr (Mm00478932_m1), Nfkb1 (Mm00476361_m1), Nfkb2 (Mm00479807_m1)* and *Rela (Mm00501346_m1).* Gene expression levels for each sample were normalized to *Hprt* or *Gapdh* expression. Mean relative gene expression was determined, and the differences calculated using the 2ΔC(t) method.

### Immunofluorescence

#### Mouse colonic section

the colon of 7-8 week-old B6J females and males was dissected out, flushed in PBS and fixed 3 hours in PFA (4% in PBS) at 4 °C, followed by overnight incubation in sucrose (30% w/v in 4% PFA) at 4 °C. Tissue was embedded in Tissue-Tek OCT Compound (Sakura, 4583) and stored at −80 °C. Tissue from 3-4 mice was cryosectioned (8*μ*m) onto the same microscope slide and washed in PBS and incubated for 1 hour in blocking solution (2.5% BSA, 5% heat-inactivated normal goat serum, 0.1% Tween-20 in PBS) at room temperature. Slides were incubated overnight in blocking solution with the following primary antibodies (1:100): rat anti-mouse SiglecF (E50-2440, 552126 BD Biosciences), Armenian hamster anti-mouse CD80 (16-10A1, 104729 Biolegend) and rabbit antimouse p-NF-KB p65 (Ser536) (93H1, 3033S Cell Signalling). After washing 3x with PBST (0.1% Tween in PBS), the following secondary antibodies were added (1:400 in blocking solution) to the slides for 1 hour at RT: AlexaFluor goat anti-rat 594 (A-11007), AlexaFluor goat-anti hamster 647 (A-21451), AlexaFluor goat anti-rabbit 488 (A-11008) all from ThermoFisher. Slides were washed 4x 5 minutes with PBST, and DAPI (D9542 Sigma, 1:1000) was added to the third washing step. Slides were mounted in Prolog Gold (P36930 Invitrogen) and imaged on a Nikon Ti2-E inverted microscope, equipped with CrestOptics X-Light v3 confocal disk unit, Lumencor Celesta lasers and Photometrics Kinetix camera.

#### Human microarrays

the microarray CO246 was obtained from Biomax.us. Deparaffinized sections were subjected to antigen retrieval in 2.4 mM sodium citrate and 1.6 mM citric acid, pH 6, for 25 minutes in a steamer. Sections were washed with PBST and blocked for 1 hour at RT in blocking buffer (5% BSA, 5% heat-inactivated normal goat serum in PBST). Slides were incubated overnight at 4 °C with the following primary antibodies (1:100, in blocking buffer): mouse anti-human MBP (BMK-13, antihuman MBP (BMK-13, MCA5751 Bio-RAD), rabbit anti-human PD-L1 (E1L3N, 13684S Cell Signalling). After washing 3x with PBST (0.1% Tween in PBS), the following secondary antibodies were added (1:400 in blocking solution) to the slides for 1 hour at RT: AlexaFluor goat anti-rat 594, AlexaFluor goat-anti hamster 647 (ThermoFisher). DAPI staining, mounting and imaging were performed as above.

### Image analysis for active-to-basal eosinophil ratio quantification

The following cores were used for quantification: A1-2,5-7 (CD); B3-6 (UC); C3-7 (healthy). Cores were chosen based on presence of colonic epithelium. ND files were imported in Imaris 9.6.0 and spots objects were created in the green (MBP) and red (PD-L1) channels separately (estimated XY Diameter = 7 *μ*m, estimated Z Diameter = 4 um, Quality Filter > 6). To quantify co-expression of PD-L1 and MBP, the distance of each spot in the green channel to the nearest spot in the red channel was computed. Green spots (eosinophils) with distance to red spots < 4 *μ*m were considered as active eosinophils (co-expressing PD-L1). Green spots with distance to red spots > 4 *μ*m were considered basal eosinophils. The active-to-basal ratio was then computed by dividing the number of active by the number of basal eosinophils in each core. For Figure 2G and 7I, the active-to-basal ratio was calculated in manually drawn regions of interest comprising the lower (basal) or upper (luminal) thirds of colon crypts.

### Histological assessment of colitis

Transversal mid-colon sections (0.5cm) were fixed overnight in buffered 10% formalin solution, followed by paraffin embedding. Sections were stained with haematoxylin/ eosin. Histopathology of the colon was scored in a blinded fashion considering four categories (each scored on a scale of 0–3): epithelial hyperplasia/damage and goblet cell depletion; leukocyte infiltration in the LP; submucosal inflammation and oedema; area of tissue affected. The final score presented (0-12) represents the sums of all categories.

### Cytospin and H&E staining

10^5^ eosinophils magnetically-enriched eosinophils (pooled from the organs of 3-4 *Il5*-tg females and males, 7-8 week-old) were resuspended in 100ul PBS and cytospun for 5 minutes at 50 g into a funnel. Slides were left air-dry overnight, stained with the Microscopy Hemacolor-Rapid staining set (1.11674.0001 Sigma) according to the manufacturer’s protocol and imaged on a Nikon Ti2-E inverted microscope.

### Single-cell RNA sequencing data analysis

#### Data pre-processing and normalization

After demultiplexing of bcl files with Bcl2fastq v2.20.0.422 (Illumina) and quality control, paired-end scRNA-seq FASTQ files were processed on the Seven Bridges Genomics platform with default parameters. Downstream analysis was conducted in *R* version 4.1.0 with package *Seurat* version 4.0.3 (Hao et al., 2021). All Seurat objects (one for each of the multiplexed samples) were merged and subjected to the same quality filtering. Cells with < 200 or > 2.500 detected genes were excluded from the analysis. After LogNormalization, the count data was scaled regressing for mitochondrial reads, and principal component analysis (PCA) was performed based on the 2,000 most variable features. Clustering and UMAP visualization were performed on the merged dataset using 50 principal components and a resolution of 0.3 for the shared nearest neighbour clustering algorithm. The clusters were annotated manually based on marker gene expression. Epithelial and mesenchymal contaminants, as well as immune cell clusters not belonging to the eosinophil-lineage were excluded from downstream analysis. A cluster high in mitochondrial genes was excluded as well. The eosinophil space was analysed by subsetting clusters expressing eosinophil markers (0, 1, 2, 3, 4 and 8). The subsetted dataset was subjected to normalization, scaling and to PCA as above. Clustering and UMAP visualization was performed using 20 principal components and a resolution of 0.3 for the shared nearest neighbour clustering algorithm.

#### Differential gene expression analysis, gene set enrichment and score computation

To extract cluster markers, *FindAllMarkers* was executed with *logfc.threshold* and *min.pct* cutoffs set to 0.25. Top-ranked genes (by logFC) were extracted for illustration. For differential gene expression, *FindMarkers* was applied with *logfc.threshold* and *min.pct* set to 0. Genes were subsequently filtered based on Bonferroni-adjusted p-value < 0.05. Scores were computed with the *AddModuleScore* function. Genes used for the scores listed in Table S2. Cell cycle scoring was performed with the CellCycleScoring algorithm from Seurat, using cell cycle-related genes from (Kowalczyk et al., 2015). For Gene Set Enrichment Analysis (GSEA), differentially expressed genes were pre-ranked in decreasing order by the negative logarithm of their p-value, multiplied for the sign of their average log fold change (in R, ‘*-log(p_val)*sign(avg_log2FC)*’). GSEA was performed on this pre-ranked list using the R package *FGSEA* (https://github.com/ctlab/fgsea/) with default parameters and the Gene Ontology Biological Process database, made accessible in R by the package *msigdbr*, https://github.com/cran/msigdbr). The results were filtered for significantly enriched gene sets (Bonferroni-adjusted p-value < 0.05).

#### Trajectory inference and trajectory alignment

Trajectory inference was performed with Monocle 2.3.6 (Qiu et al., 2017; Trapnell et al., 2014) in R version 3.6.3. After creating a Monocle object using “negbinomial.size()” distribution and lowerDetectionLimit = 0.5, the analysis was performed using Seurat’s top 2000 variable features as ordering genes. Dimensionality reduction was performed using the “DDTree” method. To visualize the eosinophil differentiation, cluster annotations were projected on the inferred trajectories. Trajectory alignment of the bone marrow-blood-colon trajectories was performed by applying dynamic time warping as performed by (Cacchiarelli et al., 2018; McFaline-Figueroa *et al.*, 2019). The steady state and *C. rodentium*-challenge trajectories were set as the reference and query, respectively. Differentially expressed genes were identifying by using a full model of ‘*y ~ pseudotime*treatment*’ and a reduced model of ‘*y ~ pseudotime’* as in (McFaline-Figueroa *et al.*, 2019)

#### Pathway and regulon activity analysis

Pathway activity was calculated across eosinophil subsets with PROGENy version 1.13.2 (Holland et al., 2020) with default parameters. Gene regulatory activity was interrogated by applying SCENIC 1.2.4 (Aibar et al., 2017) with default parameters. Briefly, after expression matrix filtering (*minCountsPerGene = 3*.01*ncol(exprMat), minSamples = ncol(exprMat)*.01*), and computing correlation, GENIE3 was applied to infer potential transcription factor targets. Coexpression networks were then calculated, regulons were created and their activity was scored in cells. Regulon activities were visualized as cluster averages using the R package ComplexHeatmap (Gu et al., 2016).

#### Integration of challenge and DSS datasets

Challenge and DSS datasets were integrated using Seurat’s anchoring-based integration method using the steady state object as reference dataset (*reference.reduction = “pca”, dims = 1:50*).

#### Cell cell interaction prediction with CellPhoneDB

Ligand-receptor interaction analysis was performed using the python package CellPhoneDB (version 2.0.0, python version 3.8.5) following instructions from the GitHub repository (https://github.com/Teichlab/cellphonedb). In brief, the annotated Seurat object of isolated LP immune cells from DSS-treated B6J mice was used to test expression of known ligand-receptor interactions from the public repository of CellPhoneDB. Gene symbols were first converted from mouse to human using the biomart R package (version 2.46.3). Mean values representing the average ligand and receptor expression of annotated clusters were calculated based on the percentage of cells expressing the gene, and the gene expression mean. To determine significance of observed means, p-values were calculated using a null distribution of means calculated for randomly permutated annotated cluster labels. An interaction was considered significant if p-values ≤ 0.05. Significant ligand receptor interaction pairs between eosinophils and CD8^+^ T cells or CD4^+^ T cells were extracted, gene symbols were converted from human to mouse, and their mean values were plotted using the plot_cpdb function from the ktplots R package (version 1.1.14) (https://github.com/zktuong/ktplots). *Plotting and statistical analysis:* Statistical analysis and visualization were performed using R version 3.6.3 or 4.1.0. Statistical significance tests were performed as described in each figure legend. Unless stated otherwise all tests were significant with Bonferroni-adjusted p value < 0.05. Plots were generated with the R package ggplot2 (Wickham, 2016) or with Prism 8.4.3 (GraphPad Software).

### Flow cytometry data analysis

#### Data analysis and plotting

Flow cytometry data analysis was performed with FlowJo software (version 10.7.1 Becton Dickinson & Company). Cell counts, relative cell frequencies or mean fluorescence intensity (MFI) were used to generate graphical plots in GraphPad Prism (version 9.1.1, GraphPad). High dimensional flow cytometry data were compensated and exported with FlowJo software (version 10) and the resulting FCS files were uploaded into Rstudio (version 4.0.3 R software environment). UMAPs were generated on stochastically selected cells from each sample and FlowSOM metaclusterings were performed for all the exported events as described previously (Brummelman et al., 2019).

#### Statistical analysis

All statistical analyses were performed with GraphPad Prism (version 9.1.1, GraphPad). Non-parametric Mann-Whitney test was used for comparing two groups, while comparisons of more than two datasets was done using one-way analysis of variance (ANOVA) with Tukey’s post-test. Differences were considered statistically significant when *P* < 0.05.

### Graphical illustrations

Experimental workflows (Figures 1A, 2G, 3E and 5A) were created using a licenced version of Biorender.com.

### Data and Code Availability

ScRNA-seq data generated during this study are deposited at the Gene Expression Omnibus under access number GSE182001. The code used in this study is available at: https://github.com/TheMoorLab/Eosino-phils_scRNASeq

---

**Figure S1:**
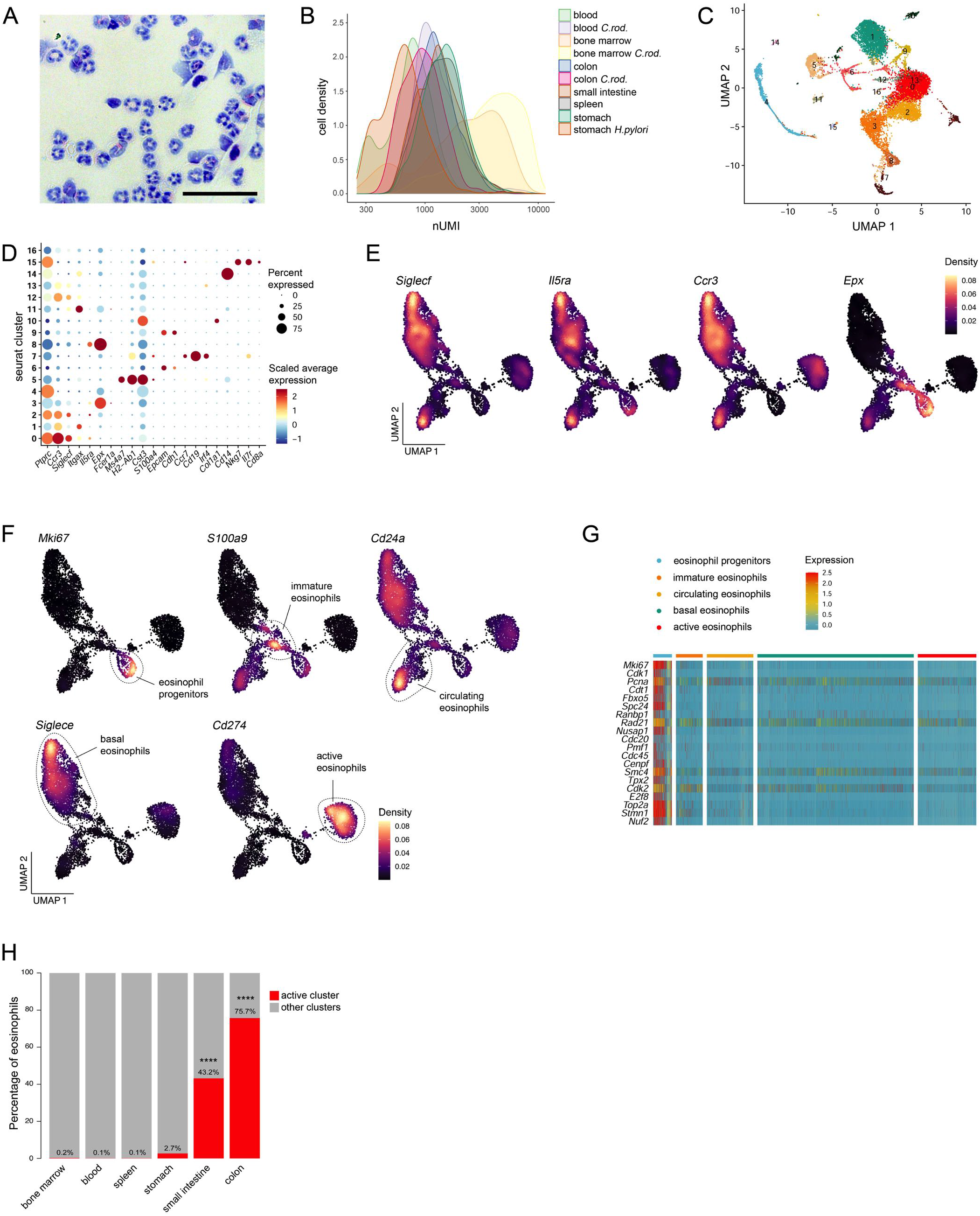
scRNA-seq reveals five distinct eosinophil subpopulations, related to Figure 1. **A)** H&E staining of eosinophils magnetically isolated from the colon of *Il5*-tg mice (n = 3). Scale bar, 50 *μ*m. **B)** Distribution of unique molecular identifiers (nUMI, log10 normalized) per cell across samples. **C)** UMAP of all sequenced single-cell transcriptomes passing quality control criteria, including contaminants not belonging to the eosinophil lineage. **D)** Expression of cluster marker genes. Dot color and size represent scaled average expression and percentage of expressing cells, respectively. Clusters 0, 1, 2, 3, 4 and 8 express eosinophil markers and were subsetted for downstream analysis. **E)** Expression density of canonical eosinophil marker genes. **F)** Expression density of subset-specific marker genes. **G)** Expression of cell cycle genes across eosinophil subsets. Rows are genes and columns are single cells, colored by scaled expression. **H)** Proportion of active eosinophils (red) across organs. P < 0.0001 (Fisher’s exact test).

**Figure S2:**
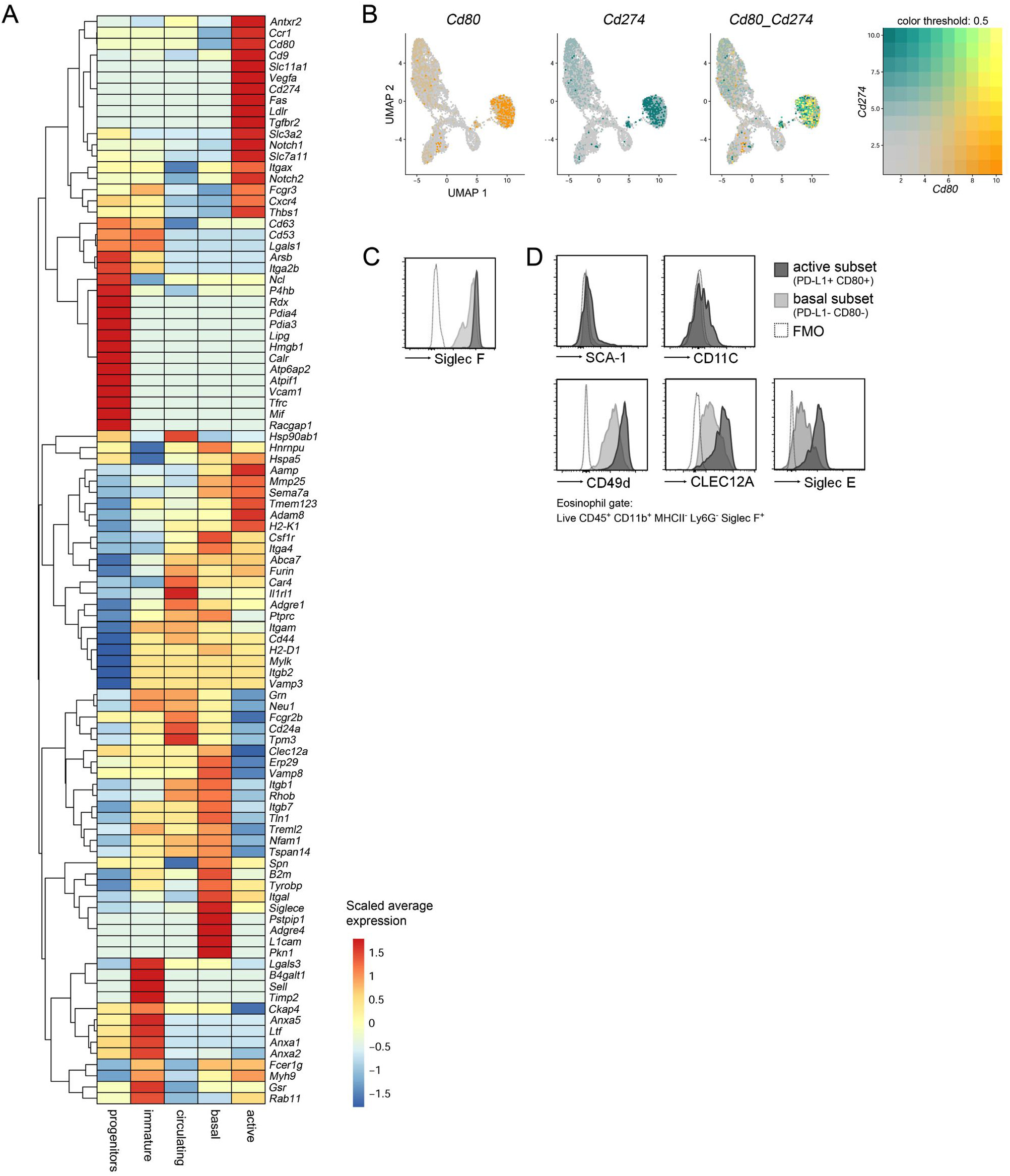
PDL1 and CD80 expression define active eosinophils in the GI tract, related to Figure 2. **A)** Scaled average expression of genes encoding surface proteins. **B)** UMAP showing co-expression of *Cd80* and *Cd274*. **C-D)** MFI of Siglec F (C) SCA-1, CD11c, CD49d, CLEC12A and Siglec E (D) in active and basal eosinophils shown in Figure 2D.

**Figure S3:**
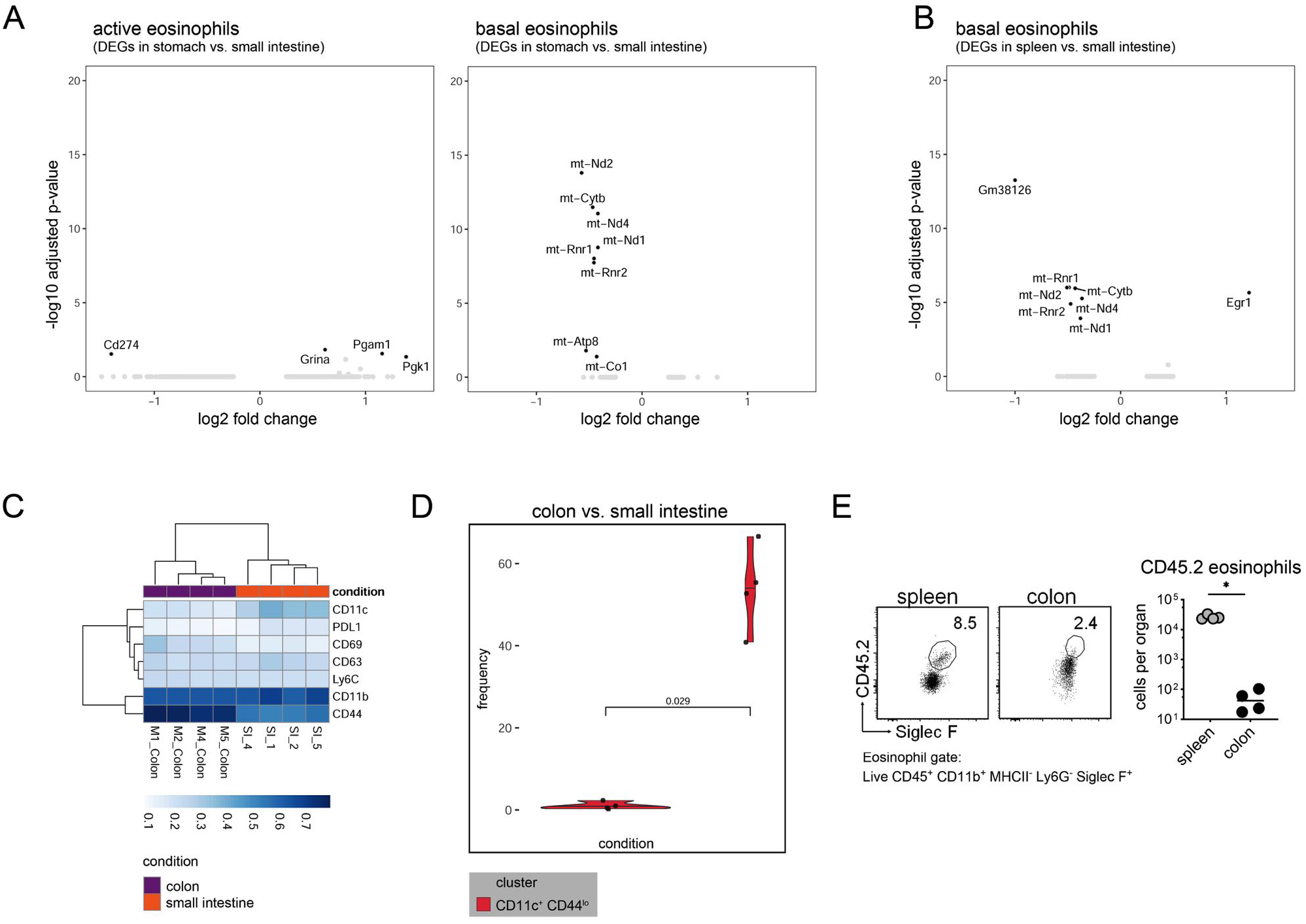
Active eosinophils display organ-specific adaptation that can be recapitulated *in vitro* and *in vivo*, related to Figure 3. **A)** Significant DEGs (average log fold-change > 0.5, adjusted P < 0.05) between colon and SI within the active (left) and basal (right) clusters. **B)** Significant DEGs between spleen and SI within the basal cluster. **C)** Heatmap showing the median expression of surface markers in colonic vs. SI eosinophils measured by high-dimensional flow cytometry (n = 4, B6J). **D)** Frequencies of CD11c^+^CD44^low^ eosinophils in colon and SI of samples shown in Figure S3D. **E)** Left: Representative flow cytometry plots of splenic and colonic eosinophils. The number of CD45.2 eosinophils as a percentage of the total eosinophils is shown. Right: absolute counts of CD45.2 eosinophils shown in Figure 3I. Medians are shown. *P < 0.05 (Mann-Whitney test).

**Figure S4:**
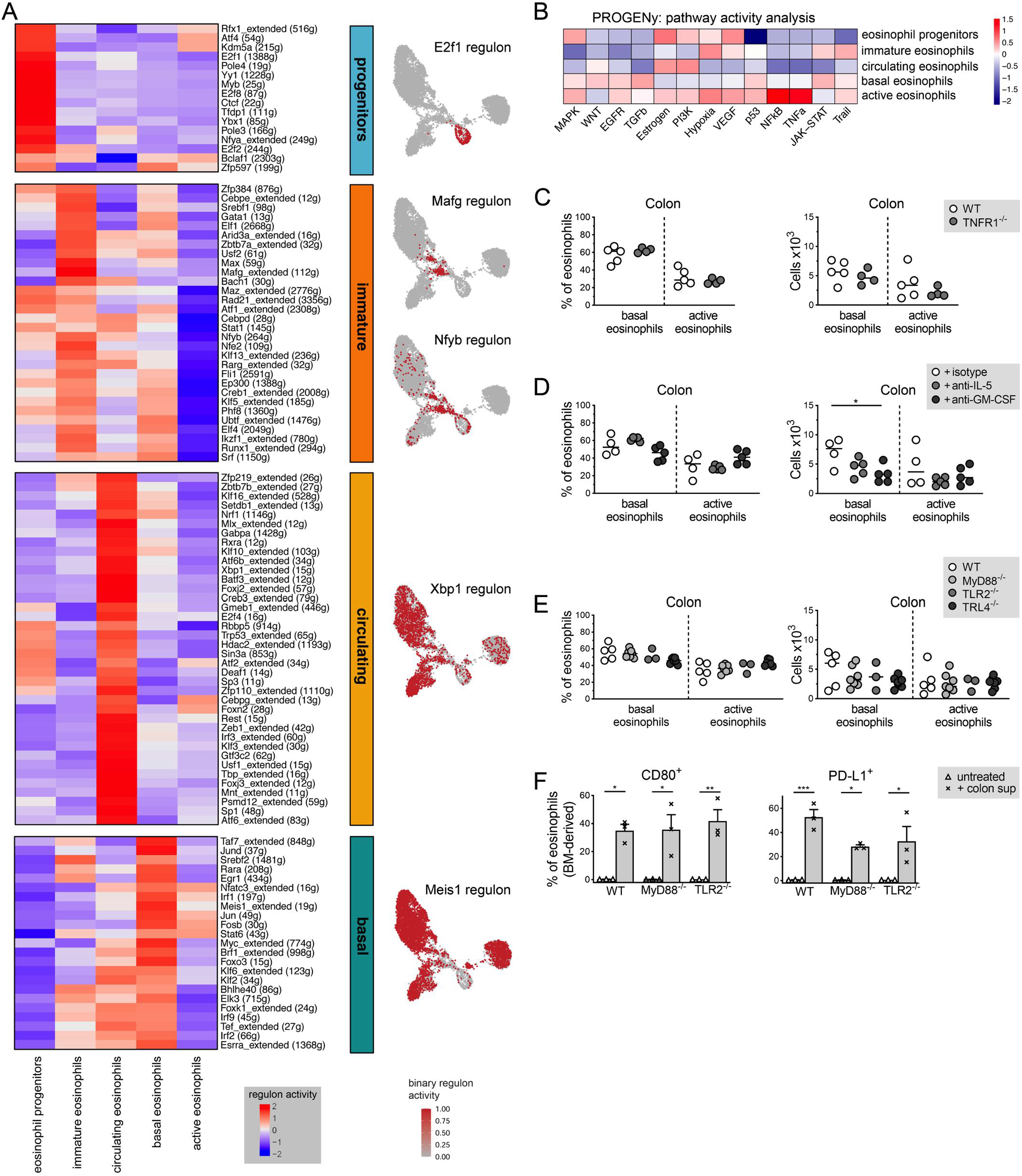
Active eosinophil differentiation is driven by NF-κB signaling and relies on bacterial cues, related to Figure 4. **A)** Regulon activity across clusters. Representative regulon projected on a UMAP plot. Cells are colored by binary regulon activity. **B)** Pathway activity across clusters according to PROGENy analysis. **C)** Frequencies and absolute counts of colonic active and basal eosinophils at steady state in TNFR1^-/-^ (n = 5) and B6J mice (n = 4). **D)** Frequencies and absolute counts of colonic active and basal eosinophils in B6J mice treated with anti-IL-5 (n = 5), anti-GM-CSF (n = 5) or isotype control (n = 4) antibodies. **E)** Frequencies and absolute counts of colonic active and basal eosinophils at steady state in B6J (n = 5), MyD88^-/-^ (n = 7), TLr2^-/-^ (n = 3) and TLR4^-/-^ (n = 7) mice. **F)** Frequencies of PD-L1^+^ and CD80^+^ BM-derived eosinophils after conditioning with colon supernatants; B6J (n = 3), MyD88^-/-^ (n = 3), TLr2^-/-^ (n = 3). Data represent the mean ± SEM. **P < 0.01; ***P < 0.001 (One-way ANOVA). C,D,E) Medians are shown. *P < 0.05 (Mann-Whitney test).

**Figure S5:**
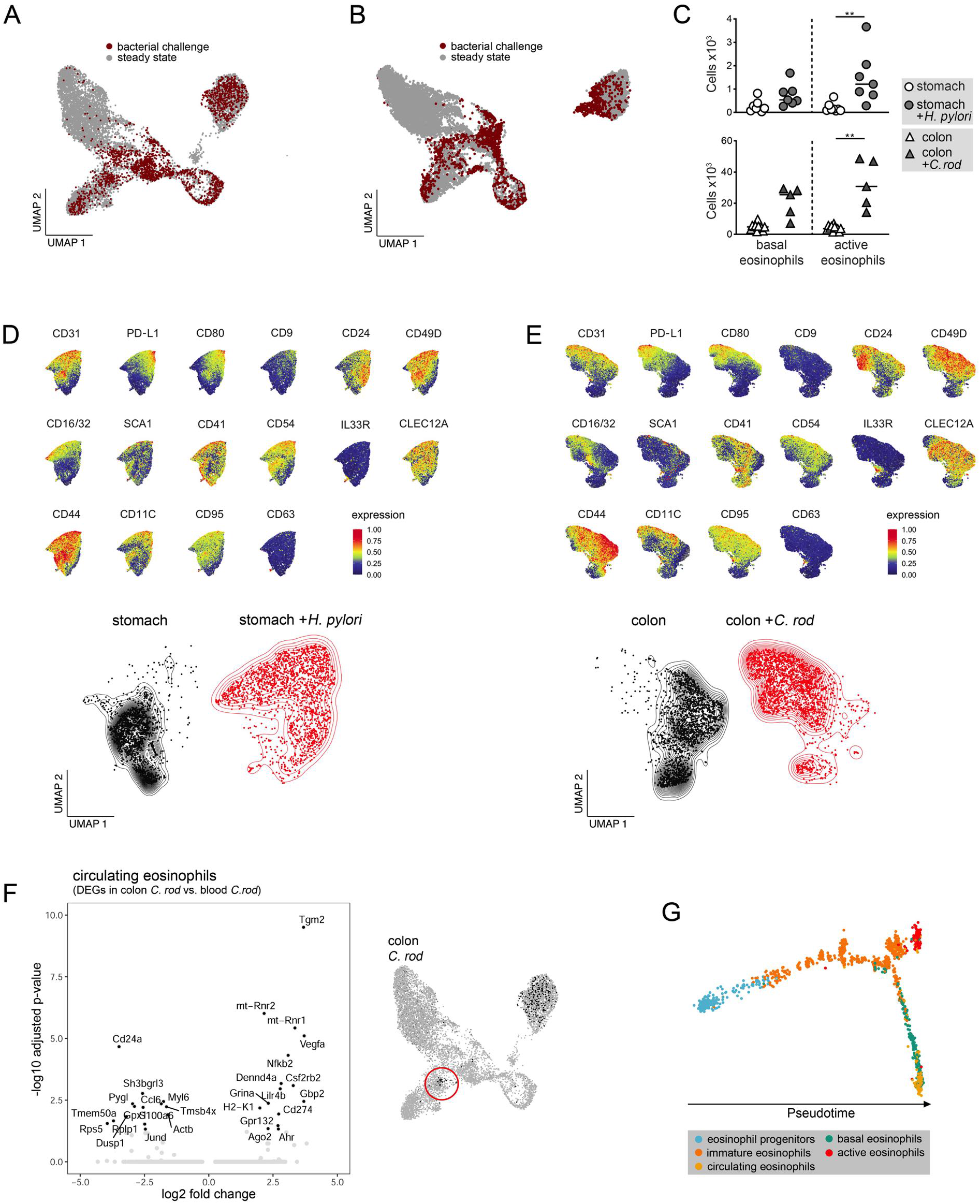
Challenge infection drives a compositional shift toward the active cluster, related to Figure 5. **A)** Data integration of eosinophils isolated from the BM, blood and colon of ***C.** rod*-infected mice, and from the stomach of *H. pylori*-infected mice (red dots). Steady state dataset (gray dots) is used as a reference. (n = 3-4, *Il5*-tg mice). **B)** UMAP of merged steady state (gray dots) and challenge (red dots) datasets. **C)** Absolute counts of active and basal eosinophils in *H. pylori*-infected stomachs (top) and ***C.** rod*-infected colons (bottom), relative to uninfected controls. Medians are shown. **P < 0.01 (Mann-Whitney test). Mice as described in Figure 5E. **D-E)** Top: UMAP showing the normalized expression intensity of 16 surface markers measured by high-dimensional flow cytometric analysis. Bottom: eosinophils in uninfected (black) and *H. pylori*-infected stomachs (D) or ***C.** rod*-infected colons (E, red) represented as split UMAPs. Mice as described in Figure 5F. **F)** Left: Significant DEGs (average log fold-change > 0.5, adjusted P < 0.05) within the circulating cluster isolated from the colon versus blood of ***C.** rod*-infected mice. Right: Red circle highlights circulating eosinophils isolated from the colon of ***C.** rod*-infected mice. **G)** BM-blood-colon eosinophil trajectory following ***C.** rod* challenge. Each dot represents a single cell colored by cluster identity.

**Figure S6.**
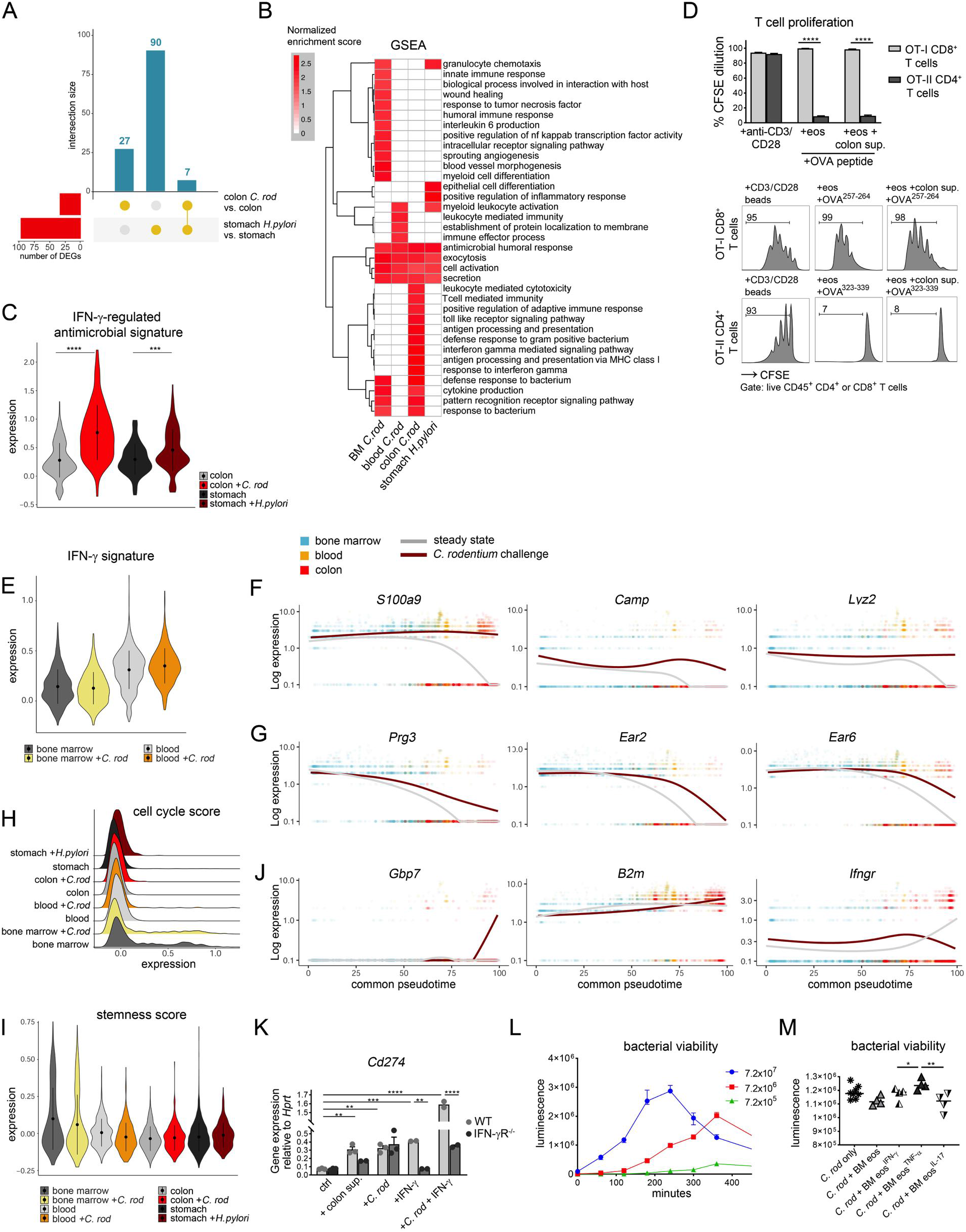
Challenge infection drives the expression of local and systemic antimicrobial gene programs, related to Figure 6. **A)** Upset plot of DEGs (***C.** rod*-infected colon vs. steady state colon and *H. pylori*-infected stomach vs. steady state stomach) within the active eosinophil subset. **B)** Significantly enriched (adjusted P < 0.05) GSEA terms within active eosinophils across challenge conditions. **C)** IFN-γ regulated antimicrobial signature across conditions. Data represent the mean ± SEM. ***P = 0.0001867; **** P < 2.2×10^−16^ (two-sided Wilcoxon test). **D)** Top: CFSE dilution of T cells co-cultured with BM-derived eosinophils conditioned as indicated and loaded with OVA peptide. Bottom: representative flow cytometry plots of the CFSE dilution. Data represent the mean ± SEM. **E)** IFN-γ signature within all clusters across conditions. Data represent the mean ± SEM. **F, G, J)** Gene expression over aligned pseudotime at steady state (gray) and upon ***C.** rod* infection (dark red). Dots indicate single cells, colored by organ: BM (blue), blood (yellow) and colon (red). **H)** Cell cycle score within all clusters across conditions. **I)** Stemness score within all clusters across conditions. Data represent the mean ± SEM. **K)** Gene expression normalized to *Hprt* measured by qRT-PCR of BM-eos upon conditioning with colon supernatant, ***C.** rod* and/or IFN-γ. B6J (n = 2–3), IFN-γR^-/-^ (n = 2–3). Data represent the mean ± SEM. **L)** *In vitro* growth dynamics of ***C.** rod* ICC180 at different colony-forming units (CFU). **M)** ***C.** rod* (ICC180) viability upon exposure for 60 minutes to BM-derived eosinophils conditioned with IFN-γ, TNF-ɑ or IL-17. Dots indicate technical replicates, eosinophils pooled from n = 2 B6J mice. In D, K, M) *P < 0.05; **P < 0.01; ***P < 0.001; ****P < 0.0001 (One-way ANOVA).

**Figure S7.**
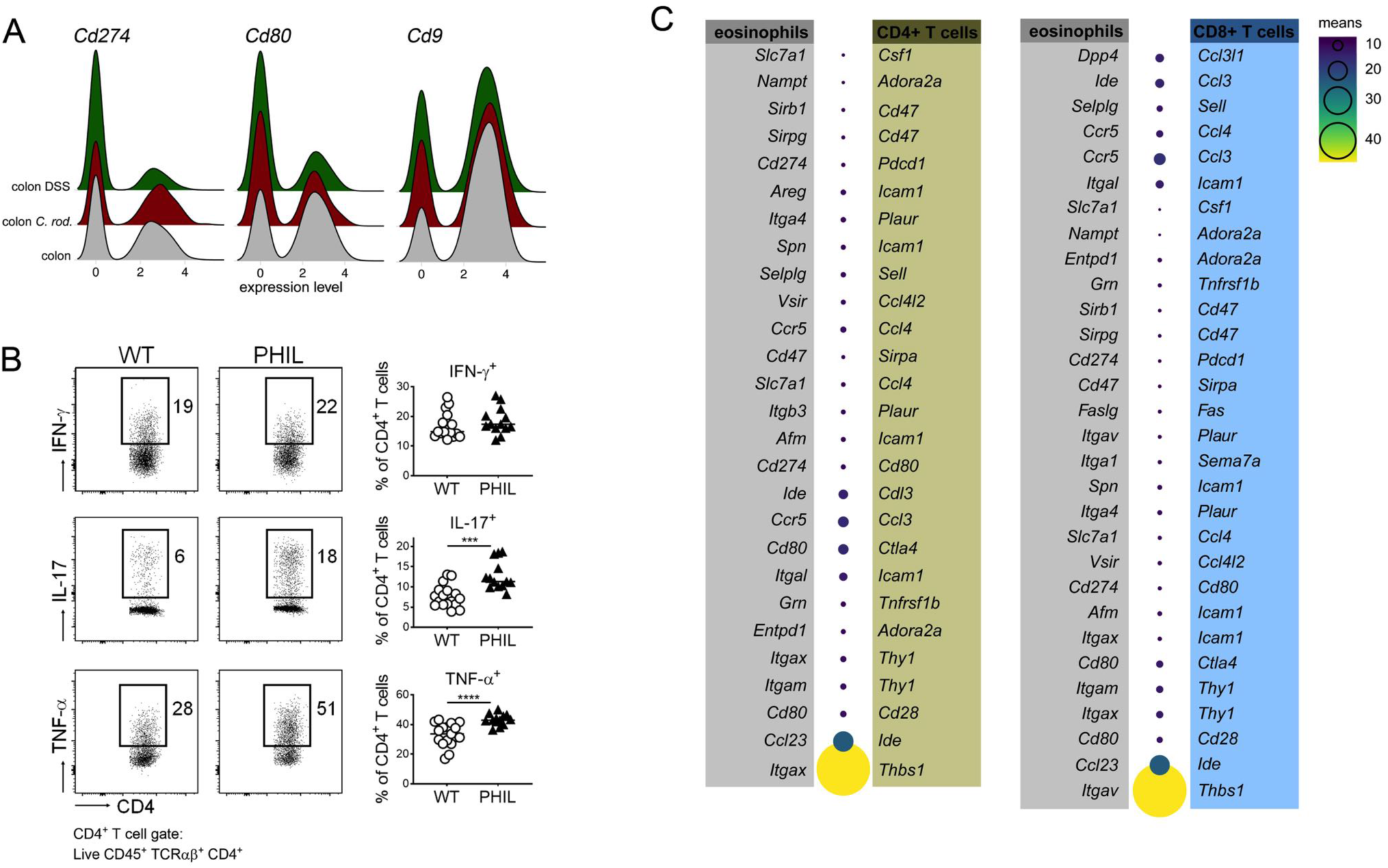
Eosinophils protect mice from experimental colitis and active eosinophils accumulate in IBD patients, related to Figure 7. **A)** Expression of *Cd274*, *Cd80* and *Cdo9* within the active eosinophil subset in the colon at steady state (gray), during ***C.** rod* infection (red) and upon DSS treatment (green). **B)** Representative flow cytometry plots (left) and frequencies (right) of IFN-γ, IL-17 and TNF-ɑ-expressing colonic CD4^+^ T cells of DSS-treated WT and PHIL mice shown in Figure 7E-F. Medians are shown. ***P < 0.001; ****P < 0.0001 (Mann-Whitney test). **C)** Ligand-receptor interactions between eosinophils and CD4^+^ T cells (left) and CD8^+^ T cells (right) predicted by CellPhoneDB. Dot size and color indicate interaction mean.

**Table S1. Differentially expressed genes in BM and blood of naïve vs. C. rod-infected mice**

**Table S2. List of genes used to define scores and signatures, with respective references.**

**Table S3. List of antibodies (antigen, clone, fluorochrome, dilution and manufacturer) used for high dimensional spectral flow cytometric analysis.**

